# Efficient genome replication in influenza A virus requires NS2 and sequence beyond the canonical promoter

**DOI:** 10.1101/2024.09.10.612348

**Authors:** Sharmada Swaminath, Marisa Mendes, Yipeng Zhang, Kaleigh A. Remick, Isabel Mejia, Melissa Güereca, Aartjan J.W. te Velthuis, Alistair B. Russell

## Abstract

Influenza A virus encodes promoters in both the sense and antisense orientations. These support the generation of new genomes, antigenomes, and mRNA transcripts. Using minimal replication assays—transfections with viral polymerase, nucleoprotein, and a genomic template—the influenza promoter sequences were identified as 13nt at the 5’ end of the viral genomic RNA (U13) and 12nt at the 3’ end (U12). Other than the fourth 3’ nucleotide, the U12 and U13 sequences are identical between all eight RNA molecules that comprise the segmented influenza genome. Despite possessing identical promoters, individual segments can exhibit different transcriptional dynamics during infection. However flu promoter sequences were defined in experiments without influenza NS2, a protein which modulates transcription and replication differentially between genomic segments. This suggests that the identity of the “complete” promoter may depend on NS2. Here we assess how internal sequences of two genomic segments, HA and PB1, may contribute to NS2-dependent replication as well as map such interactions down to individual nucleotides in PB1. We find that the expression of NS2 significantly alters sequence requirements for efficient replication beyond the identical U12 and U13 sequence, providing a mechanism for the divergent replication and transcription dynamics across the influenza A virus genome.

## Introduction

RNA viruses are constrained in genome size as a result of limitations from packaging and mutational burden.^1^ Due to these constraints it is quite common for viral genomes is to encode multifunctional proteins to maximize their use of limited genomic real estate.

Influenza A virus (IAV) is a negative-sense, segmented, single-stranded RNA virus. IAV’s eighth, and smallest, genomic segment encodes two multifunctional non-structural proteins, NS1 and NS2. NS2 is also referred to as nuclear export protein (NEP) due to its role in facilitating genome egress from the nucleus to the cytoplasm.^2,3^ This protein also aids in other functions such as viral replication and budding.^4–10^

Once IAV viral ribonucleoproteins (vRNPs) are imported into the nucleus of an infected host cell, the viral polymerase at the terminus of each vRNP interacts with host RNA polymerase II to cap-snatch and cleave a nascent transcript to act as a primer for positive-sense viral mRNA transcription.^11–14^ After viral mRNAs are translated, polymerase proteins are shuttled into the nucleus and initiate viral replication. These additional viral polymerases use the viral genomes (vRNA) as a template to synthesize positive-sense complementary RNA (cRNA), which in turn becomes a template for further production of vRNAs.^15,16^

NS2 has been shown to interact with two subunits of the heterotrimeric viral polymerase, PB2 and PB1, and alter the ratio of vRNA, cRNA, and mRNA in minimal replication assays—coordinating an increase in cRNA synthesis and decrease in mRNA transcription.^4,5,17^ Additionally, polymorphisms in NS2 can lead to variation in the frequency of defective viral genome accumulation, further supporting its role in controlling genome replication.^18,19^ Lastly, NS2 influences the response of avian influenza polymerases to ANP32A, a host dependency factor and key determinant of species tropism of avian IAV.^20–22^

Using a combination of length-variant libraries in HA and PB1, and sequence-variant libraries in PB1, we now establish that NS2 modulates IAV replication to a greater degree than perhaps previously appreciated. Most critically, we now know that the canonical viral promoters, considered to be both essential and sufficient for genome replication, cannot by themselves fully explain the dynamics of replication in the presence of NS2.^23–25^ Specifically, in our sequence-variant libraries in PB1 we identify sites that modulate viral replication in the presence of NS2 which are not conserved between IAV’s genomic segments, in contrast with the canonical viral promoter. As these sites differ between segments, they may explain how IAV fine-tunes transcription and replication between its genomic segments, which exhibit different dynamics despite sharing identical core promoter sequences.^26–30^

## Results

### Length selection differs between minimal replication assays and viral infection

Since the original development of reverse genetics there have been numerous studies of selection pressures on the genome sequence of IAV. These pressures include both broad, architectural, selection on elements that are constrained by polymerase processivity and/or nucleoprotein coating, as well as site-specific constraints such as a need to retain the minimal viral promoter and vRNP bundling sequences.^31–36^ These steps in the life cycle shape the fitness of any given viral genome and place fundamental limits on viral evolution.

These selection pressures exist alongside, and potentially in conflict with, a need to encode functional protein products. This makes it difficult to study selection pressures on genomic sequence independent of protein coding— does a mutation lead to loss of fitness because it changes amino acid sequence, or because it influences genome replication or packaging? However, this difficulty only strictly applies to replication-competent viruses. Viral populations also contain defective interfering particles (DIPs), which encode a non-standard, or defective, viral genome (nsVG, DVG).^37,38^ For IAV these genomes generally (but not exclusively) contain large internal deletions in at least one of three genomic segments encoding the heterotrimeric polymerase, PB2, PB1, and PA.^39–42^ For our purposes, DVG’s, including deletions as observed in IAV, reduce or remove selection on protein coding (other than the encoding of detrimental products), while retaining selection during genome replication and packaging.^43^ Thus these mutations can reveal selection pressures that are otherwise masked.

In a previous publication from our group (Mendes and Russell (2021)), we generated artificial length-variant barcoded, DVG vRNA libraries in the HA (hemagluttinin) and PB1 (polymerase basic 1) segments of the IAV strain A/WSN/1933.^36^ In that study, we transfected these libraries alongside plasmids encoding the minimal viral replication machinery—PB2, PB1, PA, and NP—and tracked how length of a vRNA influences replication. This work revealed that smaller variants were able to replicate much faster than their longer counterparts, and that this selection was balanced with selection for longer fragments during packaging (Fig. 1a). Therefore, at least for the lengths we tested, polymerase processivity appears to be the key rate limiting step in influenza genome replication. In contradiction to this simple model, a concurrent publication, Alnaji *et al.*, found that in a single-round viral infection competition assay, a 395nt variant of the PB2 segment was outcompeted by a full-length counterpart during genome replication.^44^

**Figure 1.**
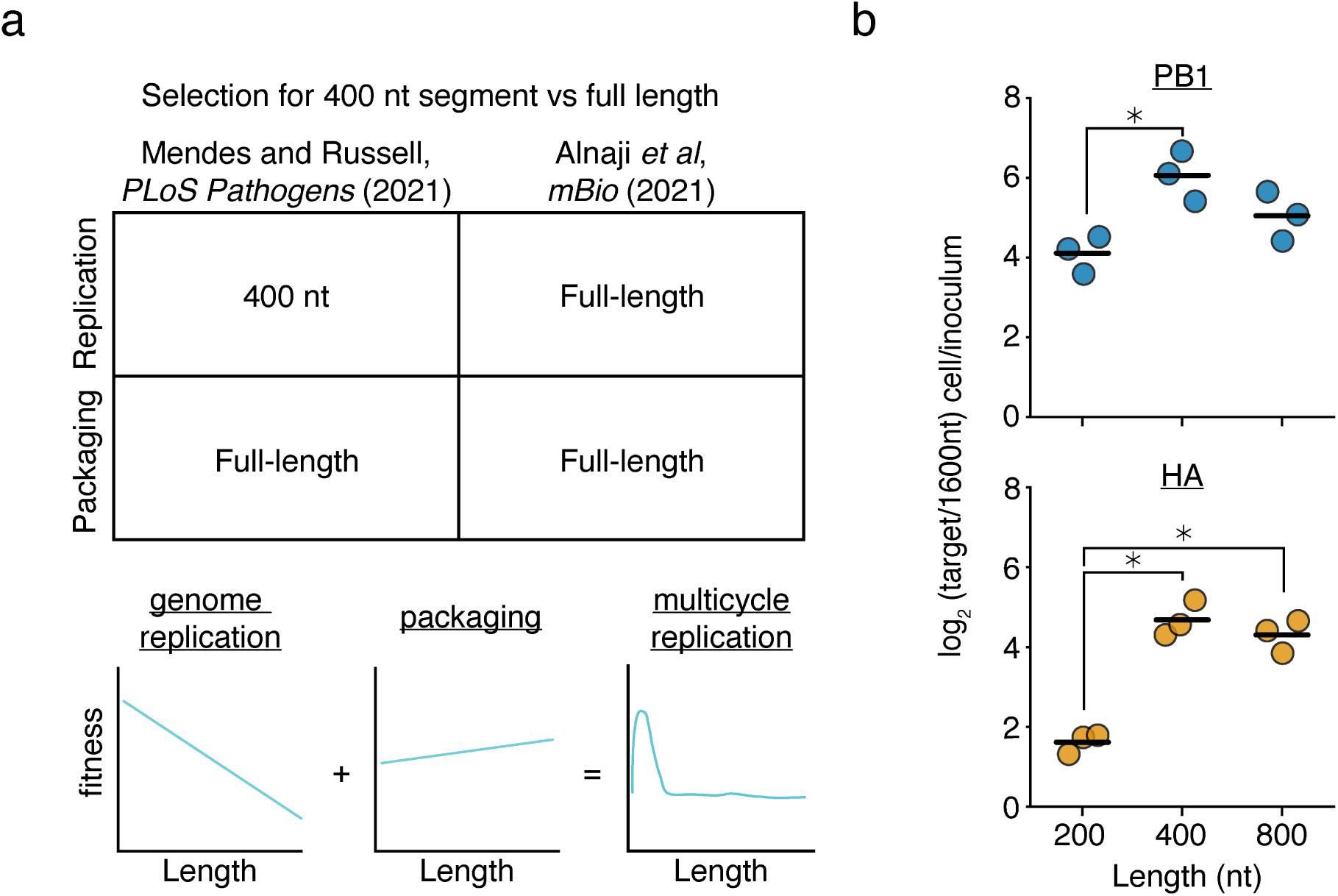
Prior models incompletely describe genome replication during viral infection. **(a)** (top)Our previous work found that a 400nt variants of the PB1 and HA segments outcompete their full-length counterparts during genome replication, but during packaging, full-length variants have an advantage. More comprehensive library-based analyses of genomic length showed that a smaller sizes correlate with better replication efficiency and longer sizes better packaging efficiency. Inconsistent with this model, Alnaji *et al.* described a phenomenon where a short variant of the PB2 segment was outcompeted during genome replication. (bottom) Schematic of our model derived from measuring thousands of length-variants in Mendes and Russell (2021).^36^ **(b)** Barcoded 200, 400, 800, and 1600nt variants of A/WSN/1933 PB1 and HA bearing equal sequence length from 5’ and 3’ ends were generated and rescued by coinfection with wild-type virus. Viral supernatant was used to infect A549 cells at an MOI of 25, and qPCR was used to analyze the proportion of each variant before infection and within infected cells at 8h post-infection. 200, 400, and 800nt variants are assessed by their frequency relative to the 1600nt species. Asterisks indicate significantly different values, ANOVA p*<*0.05 with post-hoc Tukey test, q*<*0.05. n=3, individual replicates and mean shown.

This discrepancy might be due to elements present during viral infection that are not included in minimal replication assays, or, alternatively, could be due to some effect specific to the variant studied in Alnaji *et al.*. With respect to the latter hypothesis, it has been observed that different deletions may influence polymerase processivity through the formation of alternative RNA structures, rather than, or in addition to, length-dependent effects.^45,46^ To explore further, using the same virus as our prior study (A/WSN/1933) we generated barcoded variants of PB1 and HA of 200, 400, 800, and 1600nt. We then rescued these variants using infectious virus to complement missing components, and performed single-round infection competition assays at an MOI of 25. Similar to the results from Alnaji *et al.*, and in contradiction to our simplistic model, these experiments demonstrate that there is not a straightforward, linear, relationship between segment replication and length, as in both cases a 400nt variant outcompeted its 200nt counterpart (Fig. 1b). Although for many sizes, we should note, this relationship remains intact, particularly as the least fit length we tested was 1600nt across the board. This does still mean that IAV DVGs exhibit a fitness advantage during replication owing to their length; however, the nature of that advantage is more complex than we had initially appreciated.

### Expression of NS2 influences the replication of different length segments

The differences in length-dependent replication dynamics between infection and minimal replication assays suggest there is at least one missing factor in our experiments. It has been reported that NS2 can regulate the switch from viral transcription to replication, and so this may contribute to our observations.^4–7,17^ To investigate further, we used our PB1 and HA barcoded variants in a minimal replication assay with, and without, the additional expression of NS2 (Fig. 2a). Curiously, we see different effects between the two templates; for PB1, we observe a dramatic decrease in replication of a 200nt variant relative to a 1600nt control, whereas for HA we see that all lengths tested lose some of their advantage over that same control. This effect was not unique to NS2 from A/WSN/1933, as when we repeat these experiments with NS2 from a different lab adapted H1N1 (A/PR/8/1934), a circulating H1N1 (A/Cal/07/2009), or a circulating H3N2 (A/Sydney/05/1997), we observe similar results (Supplementary Fig. 1).

**Figure 2.**
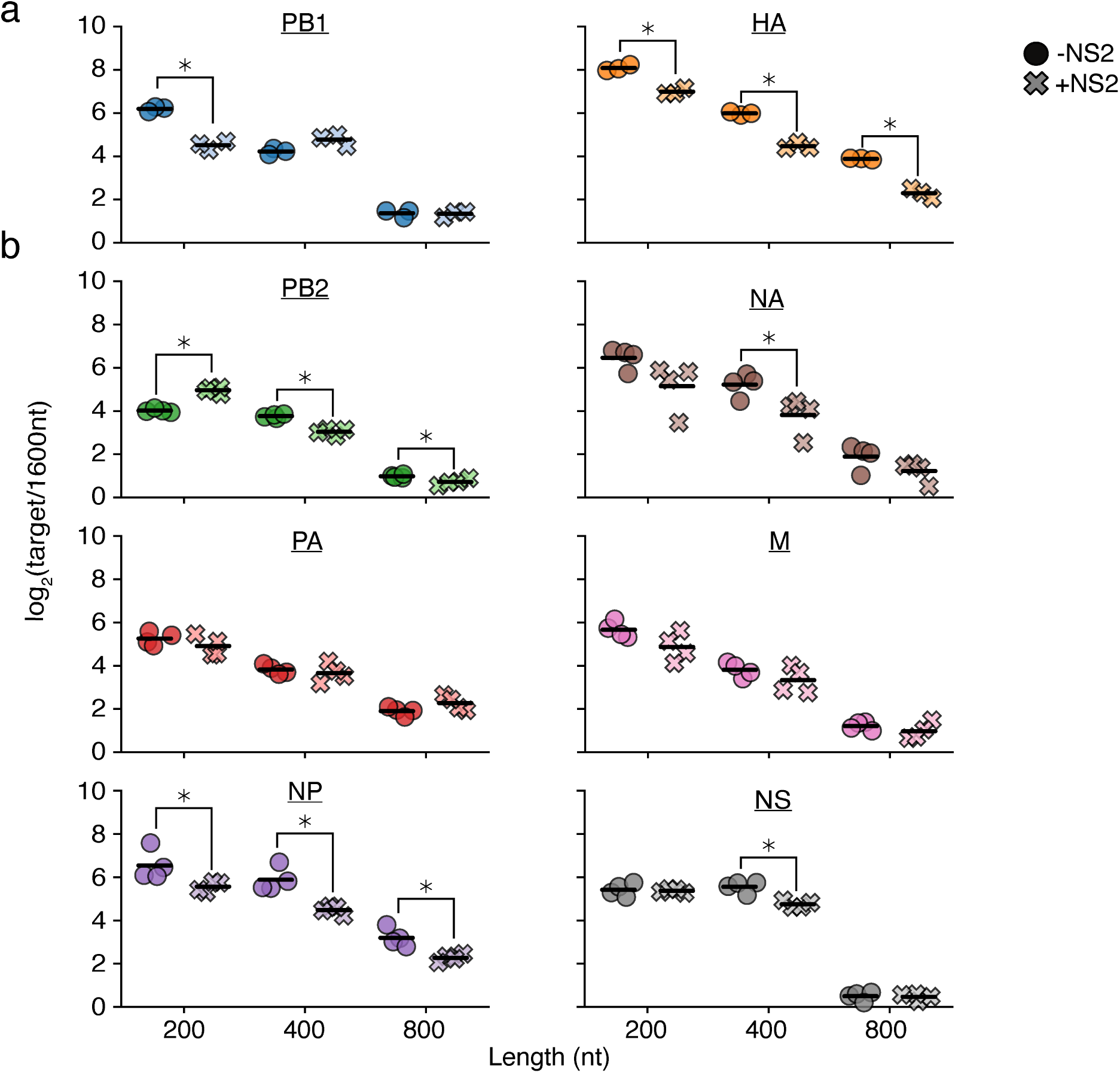
Expression of NS2 complicates simple length-dependent replication kinetics. **a,b** Barcoded 200, 400, 800, and 1600nt variants of each individual segment bearing equal sequence length derived from 5’ and 3’ ends and lacking canonical start codons were generated and co-transfected into HEK293T cells along with the minimal replication machinery with, or without, NS2. The relative frequency of each variant was measured by qPCR at 24h post-transfection, with 1600nt variants serving as a comparison. Asterisks indicate conditions significantly impacted by the expression of NS2, two-sample two-tailed t-test with a within-panel Benjamini-Hochberg corrected FDR*<*0.05. n=3 for **(a)**, n=4 for **(b)**, individual replicates and mean displayed.

The different effects between HA and PB1 prompted us to explore further. Although NS2 appears to universally promote cRNA synthesis and decrease mRNA synthesis across the eight genomic segments of IAV, the precise magnitude of these effects appears to differ between each in minimal replication assays.^4,5^ We therefore performed similar measurements across the remaining six segments (Fig. 2b). While some of these segments, such as NP, exhibited behavior similar to HA, other segments displayed trends inconsistent with either PB1 or HA. For instance, the shortest length in PB2 showed a replication advantage upon NS2 expression.

### Comprehensive analysis of the effects of NS2 expression reveals region-specific selection

Ultimately the impact of NS2 expression in minimal replication assays was highly idiosyncratic, producing a spectrum of effects between different influenza segments. One possibility for this seeming lack of pattern is that the particular junctions we chose may, or may not, be representative of the general behavior of that segment. To address this concern, and obtain more comprehensive measurements on the effect of NS2 expression on replication, we returned to our existing length-variant libraries in PB1 and HA.^36^ In brief, these libraries consist of ~1900 PB1, and ~850 HA, variants of differing sizes owing to a mixture of both deletions and duplications in every region of the segment except the first and last 75 nt, which are left unaltered as they contain the conserved influenza promoter and regions considered to be critical for packaging.

Following our prior methods, we transfected these libraries with the minimal replication machinery, either with, or without, an NS2 expression vector. The frequency of variants of any given length in the population was measured under each condition by RNAseq, and its enrichment or depletion presented in Fig. 3. We see, in agreement with with our previous results, a simple inverse relationship between segment length and replication in the absence of NS2—shorter molecular species replicate better than their longer counterparts (Fig. 3, top).^36^ Adding NS2, we see this relationship somewhat deteriorate for PB1 at the smallest lengths we test, but not for HA (Fig. 3, middle). Plotting the difference, we see that expression of NS2 dramatically reduces replication of PB1 variants ~200nt in length, and provides a slight, but consistent, selection for longer lengths in HA (Fig. 3, bottom). Both datasets closely match what we observed with individual variants by qPCR.

**Figure 3.**
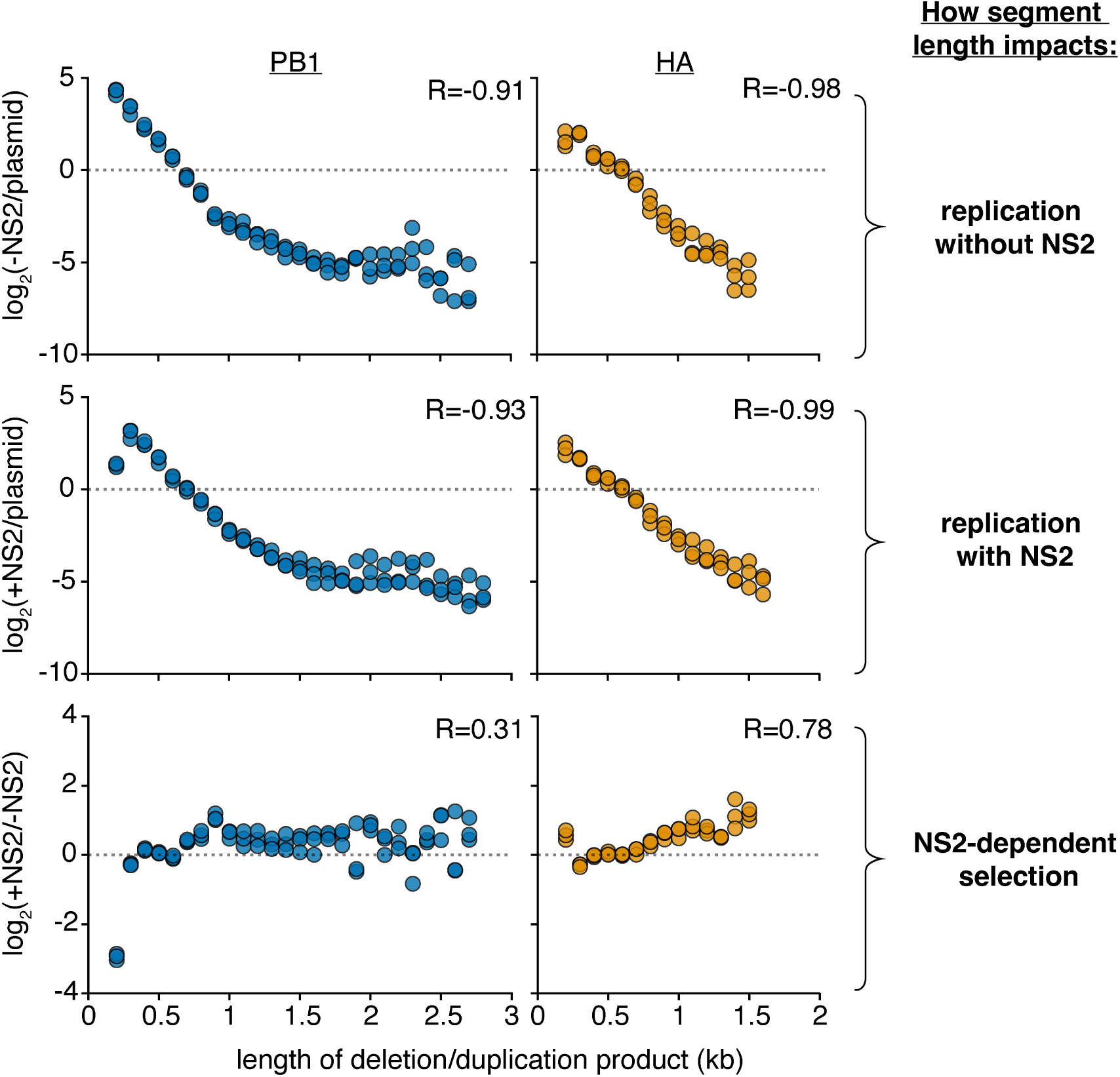
Analysis of length variant libraries confirms NS2-dependent replication effects. Size distributions of libraries in transfections with minimal replication machinery only (-NS2) and with the addition of NS2 (+NS2) 24 hours post-transfection in HEK293T cells as compared to the original plasmid library or one-another. The fraction of variants falling within each 100nt bin was compared. Points above the dotted line represent sizes in each individual library which were enriched, below, depleted. Points were only shown if represented in all three libraries under both conditions. R is the Spearman correlation coefficient. n = 3, all replicates shown. Inter-replicate correlation plots presented in Supplementary Fig. 2.

These data appear more compatible with the hypothesis that the expression of NS2 leads to sequence, or sitespecific, differences in replication rather than a length-dependent effect. Altering our analyses, we instead assess how the presence or absence of any given region in each segment impacts replication fitness, after correcting for length, presented in Fig 4a. To understand our plots, each point represents the log fitness difference as a region is *lost*. If a region is required for efficient replication, values are negative, if it is somehow inhibiting replication, values are positive.

**Figure 4.**
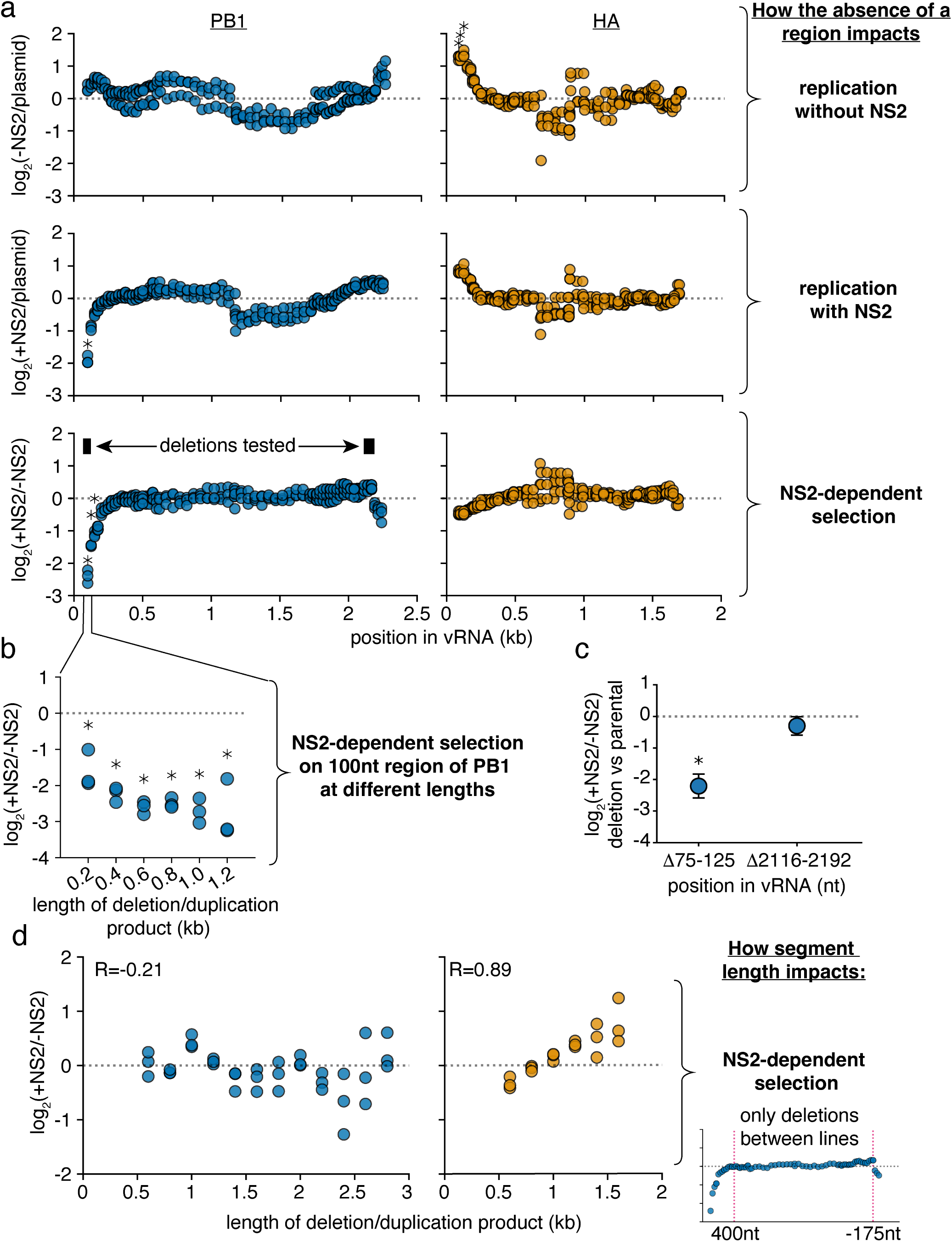
Specific regions in PB1 explain NS2-dependent replication effects. **a** Reanalysis of data from Fig. 3. The abundance of each variant that does, or does not, encode the indicated region was measured. Then, comparing only variants of identical length, the frequency at which a region is, or is not, observed under each condition was determined. Again, within only variants of identical size, changes in this frequency were measured, and the final value, presented in this graph, represents the median measurement across all lengths within a given replicate. Points above the dotted line indicate individual regions whose absence enhances replication under each indicated comparison, while points below indicate regions whose absence is associated with decreased replication. Asterisks indicate regions with a greater-than 2-fold effect size that differ significantly from no effect, one-sample two-tailed t-test, Benjamini-Hochberg corrected FDR*<*0.1. **b** The ratio of replication of sequences containing or not containing a region centering at the 100nt position, with, and without NS2 expression, across a range of lengths. Points are only shown if there were at least 100 measurements across all conditions. Statistics performed as in **(a)**, with a stricter FDR cutoff of 0.05. **c** Each indicated deletion was generated in a PB1_177:385_ background, co-transfected with a barcoded PB1 400nt internal control with, or without, NS2, and its replication was compared against the parental background at 24h post-transfection by qPCR. Asterisks indicate a ratio significantly less than 1, one-tailed one-sample t-test, p*<*0.05, indicating that the a given deletion reduces replication efficiency when NS2 is expressed. Points represent mean and standard deviation, n=3. **d** Library data analyzed as in Fig 3, bottom, excluding deletions that remove the indicated regions (first 400nt and last 175nt) in PB1 or HA.

In the absence of NS2, we do not see significant contributions of any particular region of PB1 to replication efficiency. In slight contrast, regions adjacent to the 5’ end of HA appear to negatively contribute to replication, although we do not investigate this phenomenon any further (Fig. 4a, top). As we add NS2, in HA we do not see any significant differences in replication efficiency associated with the loss of any particular genomic region. In stark contrast, we observe that regions in the 5’ end of PB1 are absolutely required for efficient replication when NS2 is expressed (Fig. 4a, middle) Plotting the differences between replication with, and without, NS2, the NS2-specific contribution of the 5’ end of PB1 to replication is quite apparent (Fig. 4a, bottom).

As sequence and length are difficult to untangle, we additionally probed whether the effects we see at the 5’ end of PB1 were true across a variety of different length variants; that is, do these sequences associate with enhanced replication no matter which length is tested (Fig. 4b)? As we would hope, when we explore our library data further, the absence of the region immediately proximal to the 5’ end of the PB1 vRNA was consistently associated with reduced replication when NS2 was expressed.

Confirming this finding, we used a 562nt PB1 template, PB1_177:385_, based on a DVG which we have previously characterized, and generated further deletions in an apparent NS2-dependent (75-125nt), and NS2-independent (2116-2192nt) region.^36,47^ This DVG consists of 177nt from the 5’, and 385nt from the 3’, end of the PB1 vRNA. We chose to use this DVG as it replicates better than full-length PB1 in minimal replication assays and so provides a broader dynamic range for our measurements. Including in each experiment a PB1 variant of 400nt as an internal control, we performed minimal replication assays with PB1_177:385_, or each individual deletion, separately. Comparing against PB1_177:385_, we find that a 5’ deletion does, specifically, impact replication when NS2 is expressed. (Fig. 4c). As we might expect of a region involved in replication, when we explore naturally occurring variation we also find that there is significantly less diversity observed in the NS2 responsive region as compared to the NS2 unresponsive (Supplementary Fig. 3).

Finally, we wished to understand whether we might be able to explain why replication of the HA segment exhibits a slight preference for longer variants when NS2 is expressed, but replication of PB1 does not. We hypothesized that the strong effect of the 5’ sequence in PB1 on NS2-dependent replication might mask the much more subtle effect we observe in HA. To test this hypothesis, we excluded any variant that removed the first 400nt, or last 175nt, from PB1 or HA, and repeated the analysis from Fig 3 to measure NS2-dependent selection. We find no evidence to support our hypothesis, as while we still see that expression of NS2 preferentially supports the replication of longer variants of HA, no such effect is observed in PB1 (Fig. 4d).

### Specific sequences are required for efficient replication in the presence of NS2

From these data it appears that NS2 expression affects replication efficiency in response to particular genomic regions or specific key sequences, rather than length. Therefore, we wanted to explore these interactions at the nucleotide-level in a single influenza segment. Also, as the first and last 75 nt of the 5’ and 3’ termini were excluded in our prior analysis, we sought to extend our understanding to include these regions. These regions contain the U12 and U13 canonical influenza promoter sequences that are essential for genome (vRNA) and antigenome (cRNA) production, as well as adjacent noncoding and coding sequences that are critical for the successful packaging of vRNPs into the virion.

To accomplish this, we performed random mutagenesis to introduce mutations into PB1_177:385_. We chose this template for two reasons. First, central positions do not appear to strongly influence replication in the presence or absence of NS2 in PB1 (Fig. 4a). By removing these positions from our analysis, we reduce the number of sequences required to appropriately sample our library, improving accuracy. Second, as in our deletion analysis, using a template that is well-replicated provides us with a greater dynamic range for our measurements.

Our PB1_177:385_ library was generated using error-prone PCR to introduce a per site mutation rate of ~0.8%. (Supplemental Fig. 4) We opted for this rate rather than the goal of only a single mutation per template to achieve variation well-above the Illumina sequencing error rate (~0.1%). We assume that the majority of sites likely exhibit no effect so it is unlikely for epistasis to significantly complicate our measurements.

We then used this library in minimal replication assays with (+NS2), and without, additional expression of NS2 (-NS2), and compared against our initial library (plasmid) or one-another (Fig. 5, Supplementary Figs. 5,7) We sequenced the RNA from these experiments using two different methods. First, we performed traditional amplicon sequencing by converting RNA to cDNA using a 3’-specific vRNA primer. Second, we performed 5’RACE (rapid amplification of cDNA ends) to analyze the first and last 20nt that are typically lost in the reverse transcription primer during amplicon-sequencing (Supplementary Fig. 8)).^48^ A consideration when exploring our data is that measurements at the first 20nt are from vRNA templates, measurements at last 20nt from cRNA templates, and measurements at the internal sites represent a mixture of signals from both cRNA and vRNA, which spontaneously convert to cDNA.^27^

**Figure 5.**
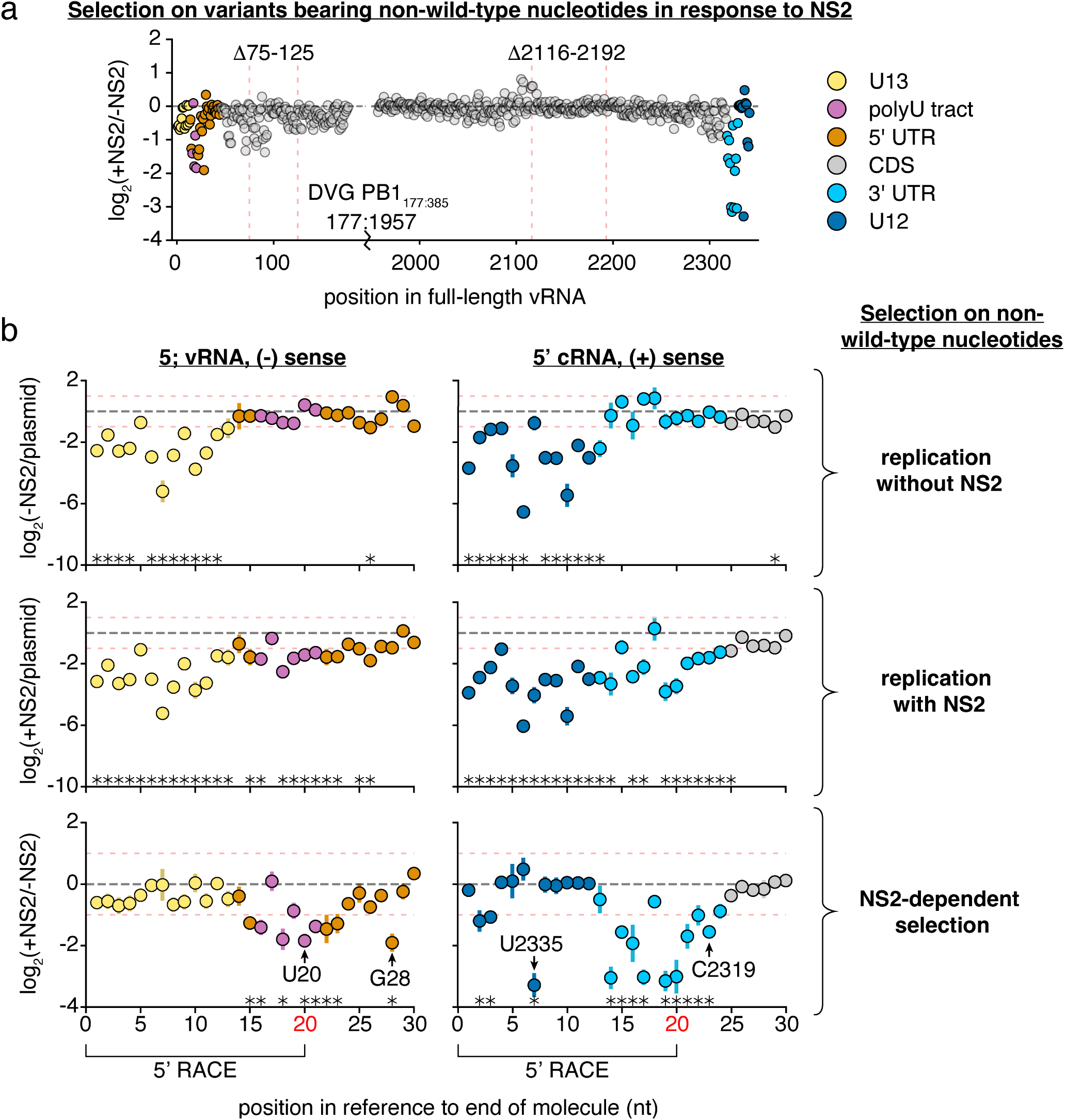
Positions within and beyond the canonical promoter are required for efficient replication in the presence of NS2. The frequency of total non-wild-type nucleotides at each position during minimal replication assays was measured under each condition, and its enrichment, or depletion shown. Similar data were procured for total information content, which considers selection at each individual nucleotide at each individual position (Supplementary Fig. 5) **a** NS2-dependent selection measured across all of PB1_177:385_ against non-wild-type nucleotides, average value across all three replicates provided. Coordinates of deletions from Fig. 4 noted. Coordinates given as the full-length vRNA, regions of functional interest annotated. Data from replication without NS2, and replication with NS2, provided in Supplementary Fig. 7. **b** Selection on non-wild-type nucleotides in the first 30nt in the vRNA (left) and cRNA (right). Mean and standard deviation graphed. Asterisks indicate regions with a greater-than 2-fold effect size (denoted by red dotted lines) that differ significantly from no effect, one-sample t-test, Benjamini-Hochberg corrected FDR *<*0.1. The first 20nt of vRNA and cRNA were inferred from 5’ RACE rather than simple amplicon sequencing (Supplementary Fig. 8). Positions further explored in Fig. 6 noted. Inter-replicate correlation plots presented in Supplementary Fig. 9.

As before, we measured how each individual variant competes with the total library during genome replication with, and without NS2. For each nucleotide position we generated two summary statistics, either measuring the collective enrichment or depletion of all non-wild-type nucleotides, or measuring the per site information content (sites that prefer only a single nucleotide, for instance, have a high information content, Fig. 5, Supplementary Fig. 5). When examining Fig. 5, positions with large negative values indicate sites where mutations tend to reduce replication, whereas positions with large positive values indicate sites where mutations away from wild-type actually increase replication. For the vast majority of sites, there is no strong preference for a wild-type or non-wild-type nucleotide at that position.

First in the absence of NS2 (-NS2/plasmid) we predominantly see strong selection against non-wild-type nucleotides within the conserved U12 and U13 promoter sequences (Supplementary Fig. 7). This is consistent with prior mapping of the viral promoter to the U12 and U13; those variants that have mutated versions of these promoters do not replicate as efficiently as wild-type.^23–25^ As we explore the effects of NS2 expression on replication, we first note that the 75-125nt region we have already defined as impacting NS2-dependent replication kinetics has several sites (82,83, and 91) which, when mutated, significantly impact replication in the presence of NS2—these, and adjacent positions, likely explain our results regarding deletions in this region (Fig. 5a, Supplementary Table 1). However, focusing on the strongest effects we observe, these tend to be more restricted to either end of the viral genome. Mutations appear to negatively impact NS2-dependent replication at sites in the cRNA/mRNA promoter and a number of other positions adjacent to both the cRNA/mRNA promoter and the vRNA promoter. We present both the first, and last, 30nt of our template to better display these positions (Fig. 5b). We see similar results in our information content analysis, which considers selection across all four possible nucleotides at each position (Supplementary Fig. 5). If sequence identity at these sites influences replication in infection, we would anticipate that they may exhibit more evolutionary constraints than other positions in the viral genome. In a natural sequence analysis we find evidence supporting this hypothesis, with significantly less variation at the 27 sites we identify as critical for efficient replication in the presence of NS2 (NS2+/NS2-) than other sites in PB1 (Supplementary Fig. 10). Curiously, despite their conservation between PB1 segments of diverse IAV strains, several of these sites are not absolutely conserved between IAV segments (Supplementary Fig. 11). This indicates that NS2-dependent effects on replication likely exhibit different requirements between different IAV segments. Altogether these findings indicate that, under polymerase activity supported by NS2, sequence identity beyond the canonical promoter influences genome replication.

We next sought to validate our findings. We chose four positions where mutations significantly effected replication in response to NS2, two that were covered by RACE sequencing, and two that were covered by amplicon sequencing (Fig. 5b). Additionally, these four positions cover a range of different functional annotations, including a position within a tract of uracils critical for stuttering and generating polyadenylated viral mRNAs.^49–51^ Into each of these positions, we introduced a transversion mutation into PB1_177:385_. We tested each mutation in minimal replication assays with, and without, NS2, co-transfected with a barcoded, 400nt, PB1 internal competitor (Fig. 6a). Our individual experiments matched our expectations from our library work, as each of these mutations in PB1_177:385_ led to a reduction of replication fitness in an NS2-dependent fashion when compared against their parental segment.

**Figure 6.**
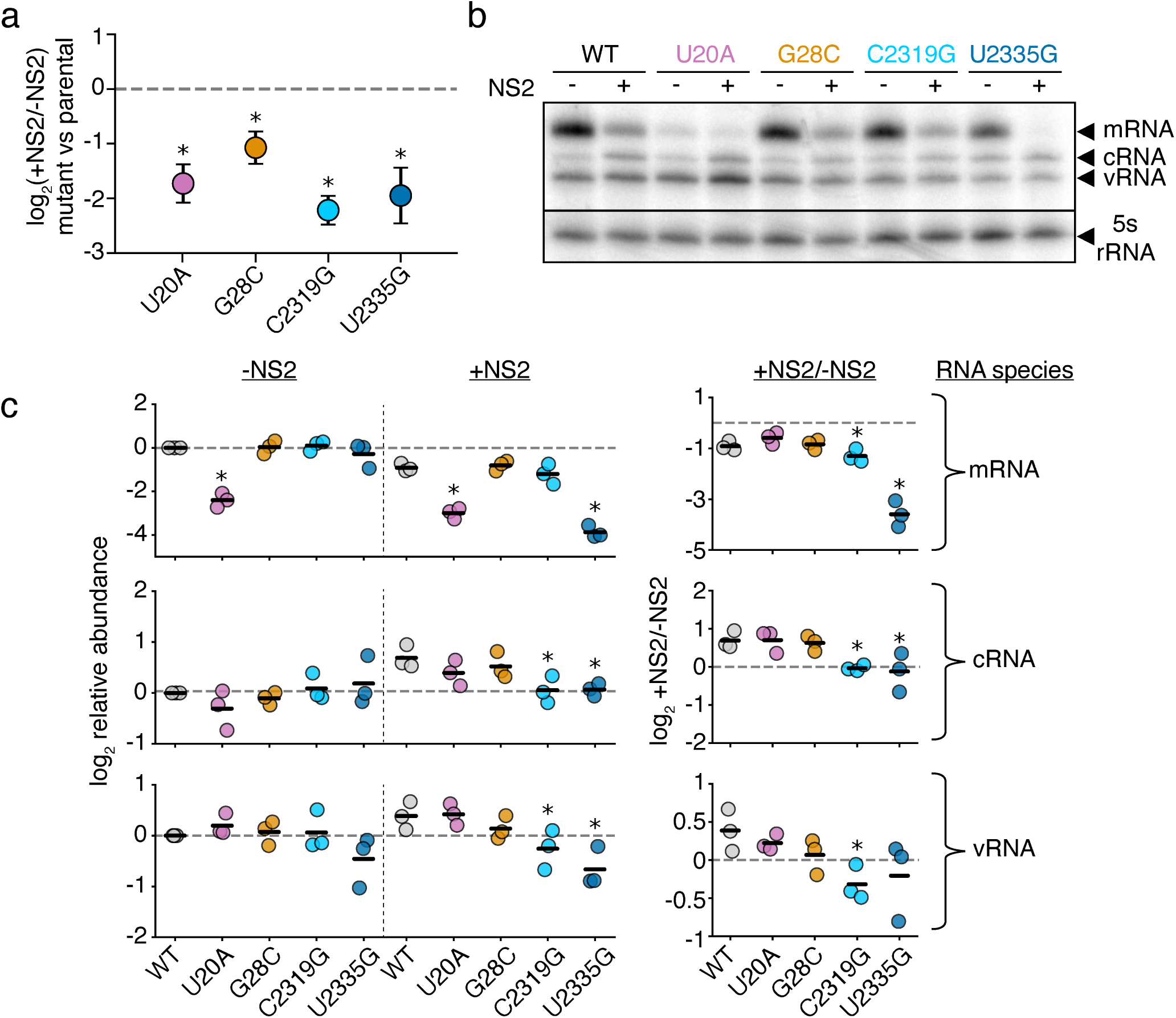
Mutations to NS2-dependent sites can influence mRNA/cRNA/vRNA ratios. **a** Each individual mutant in PB1_177:385_ was transfected alongside a 400nt internal control and minimal replication machinery with, and without, NS2. The relative frequency of each variant relative to the 400nt control was compared against wild-type under each condition. Asterisks indicate a ratio significantly less than 1, one-tailed one-sample t-test, Benjamini-Hochberg corrected FDR*<*0.05, indicating that the parental exhibits an NS2-dependent advantage over the variant. Points represent mean and standard deviation, n=3. **b** Representative primer-extension analysis of minimal replication assays with the indicated variants in PB1_177:385_ with (+) and without (-) the additional expression of NS2. Each molecular species indicated. **c** Quantitative analysis of primer extension as presented in **(b)**. All values in left two columns corrected against a parental template in the absence of NS2. Dotted line represents that value, points above indicate an increase in that molecular species, below, decrease. Values in the right column represent the ratio of points between the left two columns. Asterisks indicate values that are significantly decreased relative to the parental template, one-tailed t-test with Benjamini-Hochberg corrected FDR *<*0.1. Full gels presented in Supplementary Fig. 12. Individual replicates and mean presented, n=3. Similar analyses for full-length PB1 template for cRNA presented in Supplementary Fig.13.

With these individual mutations in-hand, we wanted to know whether we have dramatically influenced the ability of NS2 to modulate vRNA/cRNA/mRNA ratios, or whether our mutations instead have more subtle effects on replication kinetics. We tested each individual variant in minimal replication assays with, and without NS2 and measured each molecular species (mRNA/cRNA/vRNA) by primer-extension (Fig. 6b,c). Consistent with prior reports, wild-type sequence showed a reduction in relative mRNA levels and an increase in cRNAs and vRNAs with NS2 expression.^4^

As we extend our observations to our mutations, we see that all retain a reduction in mRNA levels upon NS2 expression (Fig. 6c, top). This difference is more dramatic for C2319G and U2335G, and U20A exhibits significantly reduced mRNA regardless of NS2 expression—disruption of the stuttering mechanism tied to the polyU tract could be resulting in aberrant, reduced, or absent polyA tailing and a subsequent decrease in mRNA stability.^52,53^ For cRNA and vRNA, neither 5’ mutation (U20A or G28C) produces a significant change in absolute amount or NS2-driven dynamics, at least within the sensitivity of this assay (Fig. 6c, middle, bottom). This finding may indicate that the phenotypes we observe during competition-based assays are due to differences in the relative kinetics of vRNA or cRNA production upon NS2 expression, but do not completely remove the ability of NS2 to influence replication of these templates. For C2319G, and U2335G, however, we observe that there is no apparent increase in cRNA production upon NS2 expression. Therefore these two templates appear to break the ability of NS2 to modulate the production of cRNA. To determine if these effects were unique to the template we tested, we also performed these analyses on full-length PB1 and examined cRNA (Supplementary Fig. 13). While slightly different, as our 5’ mutations actually had dramatic effects on cRNA in this assay, our results were largely similar, and differences perhaps explainable by altered replication kinetics in a long, versus a short, template.

## Discussion

Our study redefines the minimal sequence requirements for efficient replication of the IAV genome. These requirements were masked in earlier studies as they lacked NS2, a key factor promoting the replicative form of the IAV polymerase. This explanation is in accordance with two other reports that found positions that modulate replication outside of the core promoter, but did so either in the context of infection, where NS2 would be present, or in the NS segment, where NS2 would also be expressed.^54,55^ Additionally, the competitive nature of our assays, even in our reconstruction of individual variants of interest, allows us to capture kinetic differences that may not be apparent in steady-state, endpoint, assays. External to the canonical U12 and U13 promoters, we have now found that sequence identity at positions in the PB1 segment in the polyU stretch, 5’ and 3’ UTR, and even some regions within the coding sequence itself, can all modulate viral replication in the presence of the polymerase accessory factor NS2. Even within the U12 promoter, we find a single position where variants exhibit extremely different behavior during replication with, and without, NS2. These data, collectively, not only explain how NS2 can influence the spectrum of defective species observed during viral replication due to differential selection during genome replication, but also, more importantly, force us to reconsider selection across the genome during replication.

The role of NS2 in genome replication appears to be to promote the replicative, and suppress the transcriptional, form of the polymerase. A key difference in the two forms is the latter associates with the C-terminal domain of host RNA polymerase II to engage in cap-snatching of nascent transcript and acts as a single heterotrimer, whereas the former instead associates with host factors such as ANP32A and acts as a larger order oligimer (canonically thought to be a dimer).^12,14,56–59^ Explaining, in large part how NS2 accomplishes this feat, a recent cryo-EM structure of an NS2-polymerase hexamer, consisting of three dimers of NS2 and three heterotrimers of the IAV polymerase, reveals that NS2 promotes mulitmerization, and, while doing so, sterically blocks the site at which the IAV polymerase engages with host RNA polymerase II.^8^ Consistent with the physical block of transcription by NS2, while we find several sites that appear to remove the ability of NS2 to support cRNA production, none remove the ability of NS2 to decrease transcription. These are not the first mutations that can uncouple transcription and replication, for instance removal of an unpaired adenosine in the 5’ end of the vRNA within the U12 sequence decreases transcription while leaving replication intact, although they are the first that have been linked to this phenomenon of enhanced replication upon NS2 expression.^60^

Supporting our experimental work, we observe that the sites we identify as strongly influencing replication in the presence of NS2 are broadly conserved in the PB1 segment. For many of these sites, this conservation was previously noted, but attributed to packaging requirements.^61–64^ From our work, we could draw two different conclusions: 1) the first is that, as is likely true with our mutation in the polyU tract, these sites are conserved for multiple, potentially-independent, reasons, and so may be highly constrained, or 2) early measurements defining these regions as involved in packaging were often based on expression or incorporation within virions with an underlying assumption from minimal replication assays that they did not concurrently influence genome replication. As we now know that this assumption may not be correct, it may be worth re-examining some prior assumptions regarding packaging that may be better explained by an inability to efficiently replicate. We note our data by no means preclude these sites from engaging in packaging interactions, but rather provide a caution that other pressures may have produced an apparent inability to package.

Lastly, we would like to contextualize how and why NS2 might possibly engage with the sequences we identify. It is not thought that NS2 can bind to RNA directly.^2^ In its role as an export protein, it is thought to interact with M1, which interacts with nucleoprotein, which in turn interacts with the viral RNA.^10,65^ In its role in promoting replication, it has been shown to directly bind to the heterotrimeric IAV polymerase, as discussed above.^8^ What, then, explains our findings? The most parsimonious explanation which we can provide is that what we are measuring is sequence constraints on replication by the IAV polymerase in a more “true” replicative conformation. Therefore, it is not so much that NS2 directly interacts with the sites we describe, but rather the conformation the IAV polymerase adopts upon NS2 expression is more sensitive to sequence identity at these individual sites. With the recent structure of the IAV polymerase in complex with NS2, we are optimistic that our results, along with hard work from other groups, may provide a more complete, and nuanced, molecular model of IAV replication and its sequence requirements. This is of particular importance given the role of this step in defining host range, as both host cofactors and viral, including NS2, influence the ability of IAV to transition from avian to human hosts.^20^ Collectively, we hope that our data will help us all to better understand the nature and evolutionary trajectories of this important human pathogen.

## Methods

### Cell lines and viruses

The following cell lines were used in this study: HEK293T (ATCC CRL-3216), MDCK-SIAT1 (variant of the Madin Darby canine kidney cell line overexpressing SIAT1 (Sigma-Aldrich 05071502)) and A549 (human lung epithelial carcinoma cell line, ATCC CCL-185). Cell lines were tested for mycoplasma using the LookOut Mycoplasma PCR Detection Kit (Sigma-Aldrich) using JumpStart Taq DNA Polymerase (Sigma-Aldrich). All cell lines were maintained in D10 media (DMEM supplemented with 10% heat-inactivated fetal bovine serum and 2 mM L-Glutamine) in a 37*^◦^*C incubator with 5% CO2. Save where specified, all IAV experiments used A/WSN/1933, or sequences derived from the same. Genome sequence of A/WSN/1933 provided in Supplementary Data 1. Sequence of PB1_177:385_ provided in Supplementary Data 2. Sequences of NS2 from other IAV strains provided in Supplementary Data 3.

Wild-type A/WSN/1933 (H1N1) virus was created by reverse genetics using plasmids pHW181-PB2, pHW182-PB1, pHW183-PA, pHW184-HA, pHW185-NP, pHW186-NA, pHW187-M, pHW188-NS.^66^ HEK293T and MDCK-SIAT1 cells were seeded in an 8:1 coculture and transfected using BioT (Bioland Scientific, LLC) 24 hours later with equimolar reverse genetics plasmids. 24 hours post transfection, D10 media was changed to Influenza Growth Medium (IGM, Opti-MEM supplemented with 0.04% bovine serum albumin fraction V, 100 *µ*g/ml of CaCl2, and 0.01% heat-inactivated fetal bovine serum). 48 hours post-transfection, viral supernatant was collected, centrifuged at 300g for 4 minutes to remove cellular debris, and aliquoted into cryovials to be stored at -80*^◦^*C. Thawed aliquots were titered by TCID50 on MDCK-SIAT1 cells and calculated using the Reed and Muench formula.^67^ To create wild-type viral stocks with a low defective content, MDCK-SIAT1 cells were infected at an MOI of 0.01 and harvested 30 hours post-infection and titered.

### Reverse transcription and qPCR

For all qPCR experiments, cDNA was generated using the High Capacity First Strand Synthesis Kit (Applied Biosystems, 4368814). Reverse transcription was performed using the manufacturer’s protocol, save for instead of the provided primers, we used the universal vRNA primers as described in Hoffmann *et al*.^68^ Reaction conditions were an initial step of 10 minutes at 25*^◦^*C, reverse transcription at 37*^◦^*C for two hours, and an 85*^◦^*C heat inactivation step for five minutes. All qPCR experiments used Luna Universal qPCR Master Mix (NEB #M3003), with primers specified in Supplementary Data 4. qPCR conditions for all experiments were as follows; denaturation at 95*^◦^*C for one minute, then cycles of denaturation at 95*^◦^*C for 15 seconds, and annealing and extension at 60*^◦^*C fo 30 seconds. A melting curve was then performed to ensure all products are mono-peaked. Controls for each qPCR varied, for length-variant measurements the barcode associated with 1600nt variants was used as our control, for all other measurements we used a qPCR specific to a barcode associated with a 400nt variant in PB1. RNA was purified for all experiments either using the RNeasy plus mini kit (Qiagen, 74134), or the Monarch Total RNA Miniprep Kit from New England Biolabs (NEB #T2010). For measurements of cell-free viral populations, 100 *µ*l of supernatant and 600 *µ*l of lysis buffer was combined and processed.

### Generation of the mixed variant population virus and single cycle infection

Barcoded length variant segments of 200, 400, 800, and 1600nt in length in PB1 and HA were described previously.^36^ To generate viral populations of PB1 and HA containing the above-mentioned length variants, we first transfected an 8:1 co-culture of 400,000 HEK293T and 50,000 MDCK-SIAT1 cells in a 6-well plate with mixed variant populations for either PB1 and HA at equimolar ratios. In this same transfection, we included pHAGE plasmids encoding mRNA for PB2, PB1, PA, and NP downstream of a CMV promoter, also in equimolar ratios.^69^ BioT was used as our transfection reagent as per the manufacturer’s protocol. After 24 hours, D10 media was replaced with IGM and infected the cells with a low-defective wild-type WSN population at an MOI of 0.25 for 72 hours. The virus-containing supernatant was purified by centrifugation at 300g for 4 minutes. Initial viral population composition was measured by qPCR.

A549 cells were then infected with mixed populations consisting of 200nt, 400nt, 800nt, and 1600nt variants of either HA or PB1 with complementing wild-type virus at a MOI of 25, as measured by genome equivalents by qPCR. Cells were then washed to remove any additional virus at 2 hours post-infection, and, at 8 hours post-infection supernatant was removed, cellular RNA harvested, and qPCR used to measure the relative frequency of each variant. Each measurement was then corrected by the 1600nt barcode.

### Minimal replication experiments for individual variants

For all experiments, 100,000 HEK293T cells were seeded in a 24-well plate the day prior. Indicated vectors were cotransfected using BioT into our cells with equimolar amounts of plasmids encoding the minimal replication machinery (PB1, PB2, PA, and NP), with, or without, the addition of NS2. Expression constructs for minimal replication machinery for these experiments were in a pHDM backbone downstream of a CMV promoter and a *β*-globin intron. 24 hours post transfection, cells were harvested, RNA purified, and qPCR used to assess the frequency of each variant after replication.

For length-variant experiments in Fig. 2 and Supplementary Fig 1 200, 400, 800, and 1600nt variants of each A/WSN/1933 genomic segment were cloned into pHH21 for expression of authentic genomic RNA.^70^ Each of these variants possessed equal amounts of the 5’ and 3’ termini, and to each a previously-validated barcode was cloned such that they could be disambiguated by qPCR, with equivalent qPCR primers even between segments.^36^ Each variant also had the canonical start codon mutated to GTA. All clonings (including removal of start codons) used an inverse-PCR strategy and subsequent re-closing with Gibson. Primers for these clonings are provided in Supplementary Data 4. For experiments in Supplemetary Fig. 1, NS2 was cloned from each indicated strain into a pHDM expression vector. Sequences for NS2 variants provided in Supplementary Data 3.

For experiments in Fig. 4c and Fig. 6a, each variant was constructed by inverse PCR using primers described in Supplementary Data 4. Transfection of each variant was performed alongside co-transfection of a barcoded 400nt variant of PB1, as described in the prior paragraph. Ratio of each indicated variant to this control was measured under each condition, and compared against the same ratio from an experiment with the parental template. Error for these experiments was calculated using standard propagation of error.

### Mutagenesis library construction

In triplicate, a purified plasmid encoding PB1177:385 was used as a template with the GeneMorph II Random Mutagenesis kit (Agilent, #200550) in an error-prone PCR, with primers 5’-GGTCGACCTCCGAAGTTGGG-3’ and 5’-TTTTGGGCCGCCGGGTTATT-3. Following manufacturer’s instructions to achieve a 1% error rate across the 562 nt vRNA, we included 80 ng of target DNA (501.5 ng total plasmid DNA) in the mutagenic PCR reaction. Reaction conditions were 30 PCR cycles, a 54*^◦^*C annealing temperature, and 1 minute extension time. The resulting product was run on an agarose gel and purified (New England Biolab’s Monarch DNA Gel Extraction Kit, #T1020). 10 ng was subjected to an additional 10 cycles of amplification with Q5 Hot Start High-Fidelity 2X Master Mix (New England Biolabs #M0494) using the same primers as the mutagenic PCR, a 55*^◦^*C annealing temperature, and a 30 second extension time. The backbone for the mutagenic library was amplified with primers 5’-CTTCGGAGGTCGACC AGTACTCCGGTTAACTGCTAGCG-3’ and 5’-CCCGGCGGCCCAAAA TGCCGACTCGGAGCGAAAGATATAC-3’ using Q5 Hot Start High-Fidelity 2X Master Mix and reaction conditions with a 55*^◦^*C annealing temperature, a 1 minute 30 second extension time, and 28 cycle repeat. Both the mutagenic PCR and backbone were run on an agarose gel, column purified, and bead cleaned with 3X beads:sample by volume (Promega ProNex Size-Selective Purification System, #NG2001). A 20 *µ*L Gibson Assembly reaction was performed with 50 ng of vector at a 2:1 ratio for 1 hour (New England Biolabs NEBuilder HiFi DNA Assembly Master Mix, #E2621). Assembly reactions were bead cleaned with 1X beads:sample by volume to remove excess salts prior to electroporation in ElectroMax Stbl4 cells according to the manufacturer’s protocol (Invitrogen, #11635018). Cells were spread across 15 cm LB Agar + 100*µ*g/ml Ampicillin plates and allowed to grow overnight at 37*^◦^*C. Libraries were scraped and resuspended in a 10 mL LB broth. Aliquots were centrifuged until bacterial pellets appeared to be of comparable size, then plasmids were purified from these stocks (New England Biolabs Monarch Plasmid Miniprep Kit, #T1010).

### Minimal replication experiments for mutagenesis libraries

For minimal replication assays involving the length polymorphic and mutagenesis libraries, 400,000 HEK293T cells were seeded in D10 in each well of a 6-well plate 24 hours prior to transfection. Cells were transfected with helper plasmids encoding only the mRNA for A/WSN/33 PB2, PB1, PA, and NP with or without the addition of NS2 (in pHAGE vectors downstream of a CMV promoter for libraries as in Mendes and Russell, 2021), as well as the plasmid library in 1:1 ratios using BioT transfection reagent, according to the manufacturer’s protocol (Bioland Scientific LLC, # B0101).^36^ 24 hours post-transfection, cells were harvested with 300 *µ*L of RNA lysis buffer and purified using the Monarch Total RNA Miniprep Kit (New England Biolabs, #T2010).

### Sequencing preparation of mutagenesis libraries

Sequencing and analysis of length-variant libraries performed as described in Mendes and Russell, 2021.^36^ For single-nucleotide variant libraries in PB1_177:385_, for non-RACE sequencing we first converted RNA from our experiments to cDNA with a universal vRNA primer with adapter from Hoffmann *et al.*, 5’-TATTGGTCTCAGGGAGCGAAAGCAGG 3’ using Superscript III according to the manufacturer’s protocol (Invitrogen, #18080400).^68^ PCRs were performed to amplify and append partial Illumina i5 and i7 adapter sequences across four sub-amplifications of each library to sequence the entire vRNA genome, except the terminal ends which cannot be captured with this method (see 5’ RACE sequencing methods). For all reactions, 1 *µ*L of cDNA or 10 ng of plasmid were used as a template for each mutagenesis library sample was used with Q5 Hot Start High-Fidelity 2X Master Mix. The first PCR used universal primers for the PB1 segment from Hoffmann et al (5’-Bm-PB1-1-G4: 5’-TATTGGTCTCAGGGAGCGAAAGCAGGCA -3’; 3’-Bm-PB1-2341R: 5’-ATATGGTCTCGTATTAGTAGAAACAAGGCATTT-3’) with a 55*^◦^*C annealing temperature and 20 second extension time for 13 cycles, and was subsequently run on an agarose gel, extracted, and column purified. The second PCR appended Illumina intermediate sequences to staggered sub-amplifications (sequences provided in Supplementary Data 4) of each library with a 60*^◦^*C annealing temperature and 20 second extension time for 7 cycles. Sub-amplifications were pooled and purified with 3X beads:sample by volume.

To identify the sequences of the terminal ends of the vRNA and cRNA promoter that are lost during typical RNAseq amplification methods, the mutagenesis library samples were reverse transcribed with the Template Switching RT Enzyme Mix (New England Biolabs, #M0466) according to the manufacturer’s protocol and subsequently amplified and appended with the partial Illumina i5 and i7 adapter sequences using Q5 Hot Start High-Fidelity 2X Master Mix. In brief, 4 *µ*L of sample were annealed with the universal Hoffmann et al. primers for PB1 (5’-Bm-PB1-1-G4: 5’-TATTGGTCTCAGGGAGCGAAAGCAGGCA -3’ for U12 priming; 5’-TATTGGTCTCAGGGAGTAGAAACAAGGC 3’for U13 priming) and then reverse transcribed with the template switching oligo (TSO, 5’-Biotin-AAGCAGTGGTATCAACGCAGAGTACATrNrG+G-3’, containing mixed ribonucleotides at the 3rd from final position, a locked nucleic acid at the final position, and a 5’ biotin modification to prevent spurious additional concatemerization) similar to that from Picelli *et al.*^71,72^ Amplification of the 5’ terminal ends of the vRNA and cRNA strands were performed in separate reactions using a TSO-specific amplification primer (5’-AAGCAGTGGTATCAACGCAGAGT-3’) and either a vRNA amplification primer (5’-CACCATGGATACTGTCAACAGG-3’) or cRNA amplification primer (5’-TCTGAGCTCTTCAATGGTGG-3’). Amplicons were run on an agarose gel, extracted, and column purified prior to appending partial Illumina i5 and i7 adapters (common TSO partial adapter: 5’-GTCTCGTGGGCTCGGAGATGTGTATAAGAGACAG AAGCAGTGGTATCAACGCAGAGT-3’; vRNA partial adapter: 5’-TCGTCGGCAGCGTCAGATGTGTATAAGAGACAG TCCAGAGCCCGAATTGATGC-3’; cRNA partial adapter: 5’-TCGTCGGCAGCGTCAGATGTGTATAAGAGACAG CCTGTTGACAGTATCCATGGTG-3’). PCRs were run with a 65*^◦^*C annealing temperature and 20 second extension time for 15 cycles and were subsequently run on an agarose gel, extracted, and column purified.

To append final Illumina indices, a final PCR using Q5 Hot Start High-Fidelity 2X Master Mix was performed with a 62*^◦^*C annealing temperature and 20 second extension time for 7 cycles. Indexing primers were from IDT for Illumina DNA/RNA UD indexes, Set A, # 20026121. To ensure uniform amplification, samples were run on an agarose gel, extracted, column purified, and bead purified with 3X beads:sample by volume. Finally, sample purity and concentration were assessed via nanodrop and Qubit, respectively.

### Computational analysis

Assignment of barcodes in length-variant libraries was performed as described previously.^36^ For single nucleotide variant libraries, amplicon and 5’ RACE analyses proceeded with slightly different parameters. First, for amplicon analysis reads were first trimmed using Trimmomatic with the following parameters: 2 seed mismatches, a palindrome clip threshold of 30, a simple clip threshold of 10, a minimum adapter length of 2, keep both reads, a lead of 20, a sliding window from 4 to 15, and a minimum retained length of 36.^73^ Reads were then mapped against PB1_177:385_ using STAR, with relaxed mismatch paremeters (permitting a mismatch frequency of 10% over the length of the read), requiring end-to-end mapping, and an intron of 1 to remove splice-aware alignment.^74^ Nucleotide identity per site was then assessed using a custom script and a quality score cutoff of 30 (that is, sites below this score were not considered). Indels, while they are generated by the mutagenic PCR, were excluded from our analysis.

For 5’ RACE data, a number of reads were found to possess 5’ heterogeneous caps, likely either mRNA or capped cRNA sequences. These needed to be excluded such that identical molecular processes were being studied. For RNA sequencing, we enforced a match to either the sequence 5’-AAGCAGTGGTATCAACGCAGAGTACATGGGAGCGAAAGCAGGCAAACCAT-3’ or 5’-AAGCAGTGGTATCAACGCAGAGTACATGGGAGTAGAAACAAGGCATTTTT-3’ containing no more than 5 differences, as these sequences span the TSO and 5’ end of vRNA/cRNA. To produce a comprable plasmid dataset we used the sequences 5’-AAGCAGTGGTATCAACGCAGAGTCGGAGTACTGGTC GACCTCCGAAGTTGGGGGGGAGCGAAAGCAGGCAAACCAT-3’, and 5’ AAGCAGTGGTATCAACGCAGAGTCTCCGAGTCGGCATTTT GGGCCGCCGGGTTATTAGTAGAAACAAGGCATTTTT-3’, to include the plasmid backbone and permitted only 4 differences as there is a variable nucleotide in the TSO that is unaccounted for in the plasmid data.

After filtering our reads in this manner, we used STAR to map against a custom reference consisting of PB1_177:385_ with TSO adapters. Individual positions were then assessed as with our amplicon data. The first 20nt of our dataset were derived from 5’ RACE, remaining positions derived from amplicon sequencing.

For assessments of entropy and generation of sequence motifs for our mutational library, we used a calculation based on Shannon entropy with information content in bits as descibed by Schneider *et al.*^75,76^ This calculation consists of taking the total possible information content for a nucleotide position (2 bits) and subtracting the Shannon entropy. The latter is calculated by taking the sum of frequency that each nucleotide was observed multiplied by the log_2_ of that same frequency. However, instead of the frequency at which a base is observed, we scaled a presumed frequency of a nucleotide by the observed selection (that is we summed all selections, and took the relative ratio between any given selection and this summed selection). For instance, if there is no strong selection, and all nucleotides are observed at equal frequency before, and after, a given experiment, then each nucleotide is given a frequency of 25%. This would maximize Shannon entropy (2 -(0.25x-4)x 4, or -2), giving an information content of 0. Conversely, if other nucleotides were completely purged during selection, this would give us maximal information content of 2. Letter heights in sequence logos were procured by multiplying the contribution of a given selection-derived frequency of a given nucleotide by the total information content at a position. Sequence logo graphs generated by the package dmslogo (https://github.com/jbloomlab/dmslogo).

Lastly, for natural sequence diversity, we procured PB1 sequences from the NCBI flu database https://www.ncbi.nlm.nih.gov/genomes/FLU/Database/nph-select.cgi?go=genomeset on July 25th, 2024 that met the following prerequisites: full-length, 2341nt long, and exclude vaccine and laboratory strains. These sequences were then clustered by CD-HIT to an identity of 99%, and a single example of each cluster retained.^77^ Thereafter, sequences with ambiguities or indels relative to other PB1 sequences, were manually removed. This resulted in 1939, highquality, PB1 sequences that exhibit good alignment when simply lined-up with one-another, giving high confidence in the comparability of any given position, and, given the CD-HIT clustering, are unlikely to be aggressively weighted towards any single outbreak (Supplemental Data 5) At this point, no further alignment was required, and analysis proceeded.

All code was run on the San Diego supercomputer center, Triton shared computing cluster.^78^

All computational analysis presented in https://github.com/Russell-laboratory/NS2_sequence_specificity.

### Primer extension analysis

For all experiments in Fig. 6b,c and Supplementary Figs. 13 and 14, 100,000 HEK293T cells were seeded in a 24-well plate the day prior. Indicated PB1 variants in a pHH21 vector were co-transfected using BioT into our cells with equimolar amounts of plasmids encoding the minimal replication machinery (PB1, PB2, PA, and NP), with, or without, the addition of NS2. Expression constructs for minimal replication machinery for these experiments were in a pHDM backbone downstream of a CMV promoter and a *β*-globin intron. 24 hours post transfection, cells were harvested, RNA purified, and qPCR used to assess the frequency of each variant after replication. Unlike in qPCR experiments, only the indicated PB1 variant was transfected, with no additional internal control.

For RNA extraction, total RNA was extracted from cells lysed in Trizol using chlorophorm extraction and isopropranol precipitation as described previously.^79^ Approximately 400 *µ*g of RNA was subsequently reversed transcribed using 32P-labeled PB1 c/mRNA-specific primer (PB1 146: 5’-TCCATGGTGTATCCTGTTCC-3’), 32P-labeled PB1 vRNA-specific primer (PB1 2203: 5’-CGAATTGATTTCGAATCTGG-3’), 32P-labeled 5S rRNA-specific primer (5S100: 5’-TCCCAGGCGGTCTCCCATCC -3’), and a SuperScript III reaction mix (Invitrogen #18080044) containing 1 U/*µ*l RNase inhibitor (ApexBio #K1046). cRNA products were resolved using 7M Urea/12% PAGE in 1x TBE. Radioactive signals were collected using BAS-MS phosphorimaging plates (FujiFilm #28956475) and visualized using a Typhoon scanner. Densitometry analysis was performed using ImageJ.

## Data availability

Processed files and raw sequencing files can be found in the NCBI gene expression omnibus under accession GSE276697, or additionally under bioproject PRJNA1158716. Analysis code at https://github.com/Russell-laboratory/ NS2_sequence_specificity.

## Acknowledgments

This work was in part funded by the NIGMS of the NIH under grant R35GM147031 awarded to ABR. and by NIH grant DP2 AI175474 awarded to AJWtV. KAR was supported by NIH training grant 1T32GM148739-01A1 and an NSF GRFP under grant number DGE-2039656. The funders had no role in study design, data collection and analysis, decision to publish, or preparation of this manuscript. We thank Rommie Amaro and Alma Castaneda for insightful discussions regarding IAV polymerase structure.

## Supplementary Figures, Tables, and Files

**Supplementary Figure 1.**
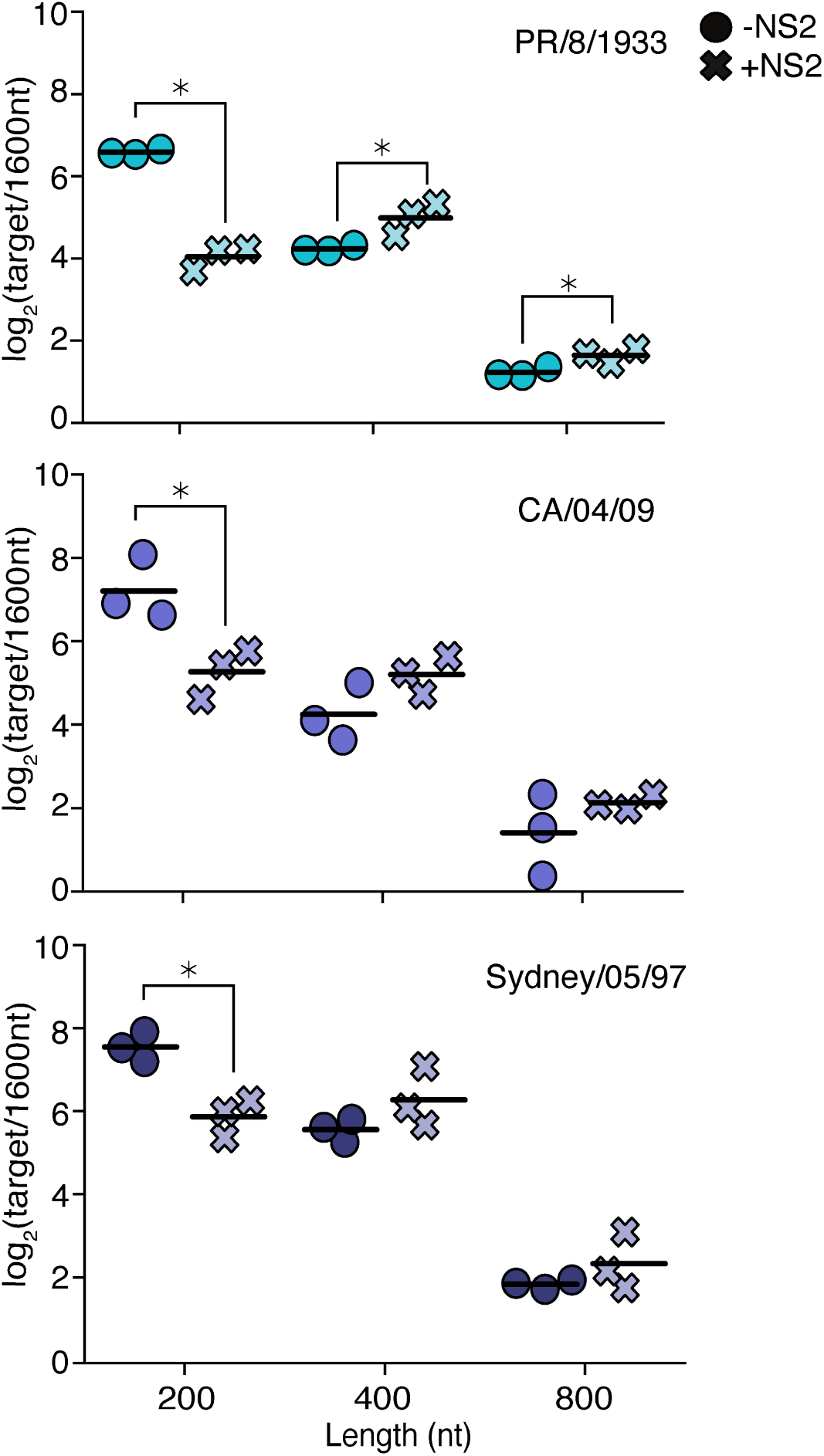
NS2 expression from diverse IAV strains exhibit similar effects on genome replication. Experiments performed as in Fig 2, using the PB1 segment and minimal replication machinery from A/WSN/1933 and NS2 from the indicated IAV strains. Asterisks indicate conditions significantly impacted by the expression of NS2, two-sample two-tailed t-test with a within-panel Benjamini-Hochberg corrected FDR*<*0.05. n=3, individual replicates and mean displayed.

**Supplementary Figure 2.**
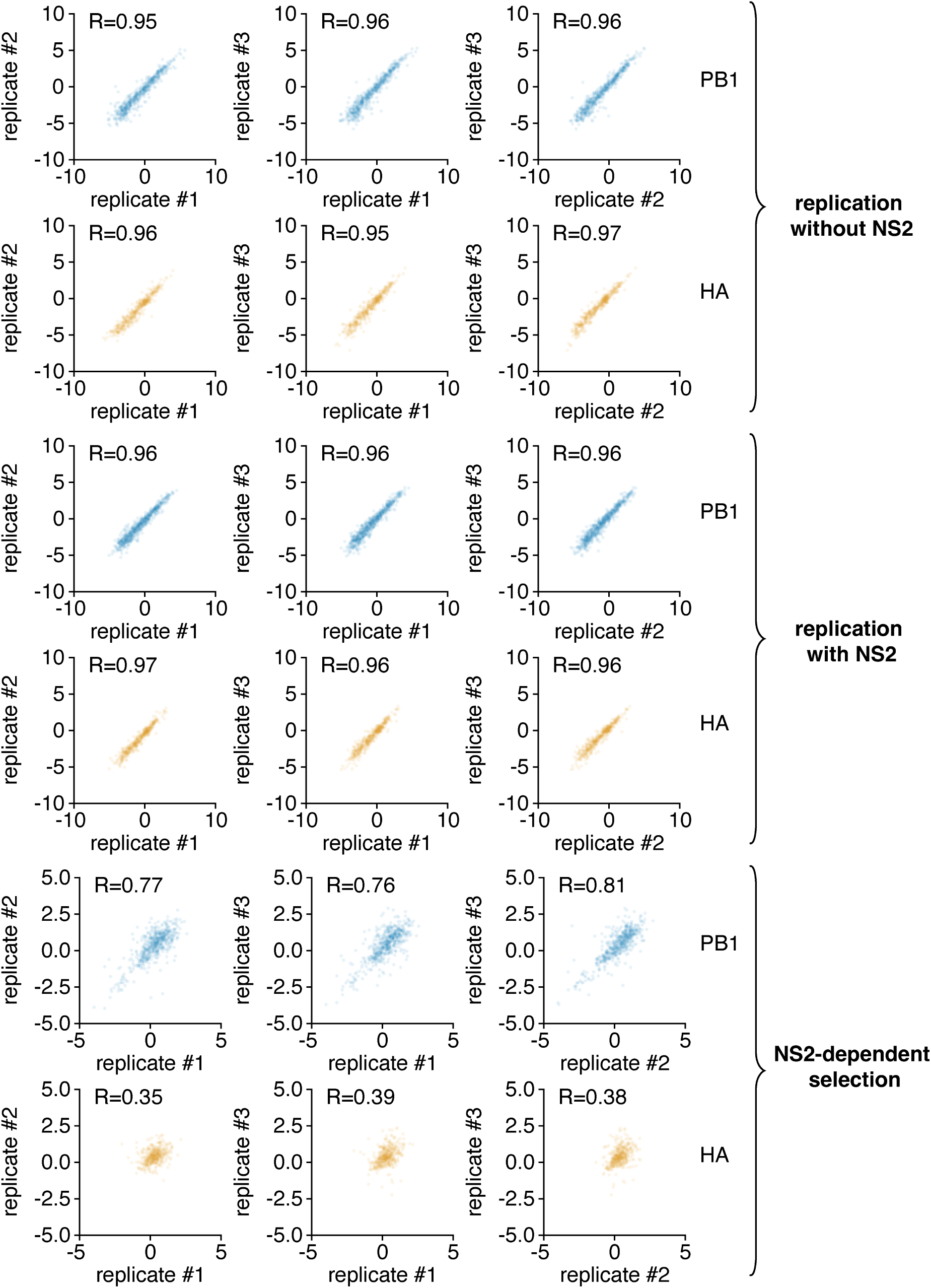
Inter-replicate correlation from Fig. 3. Inter-replicate enrichment or depletion values as calculated in Fig 3. Values are in log_2_. R is the Pearson correlation coefficient.

**Supplementary Figure 3.**
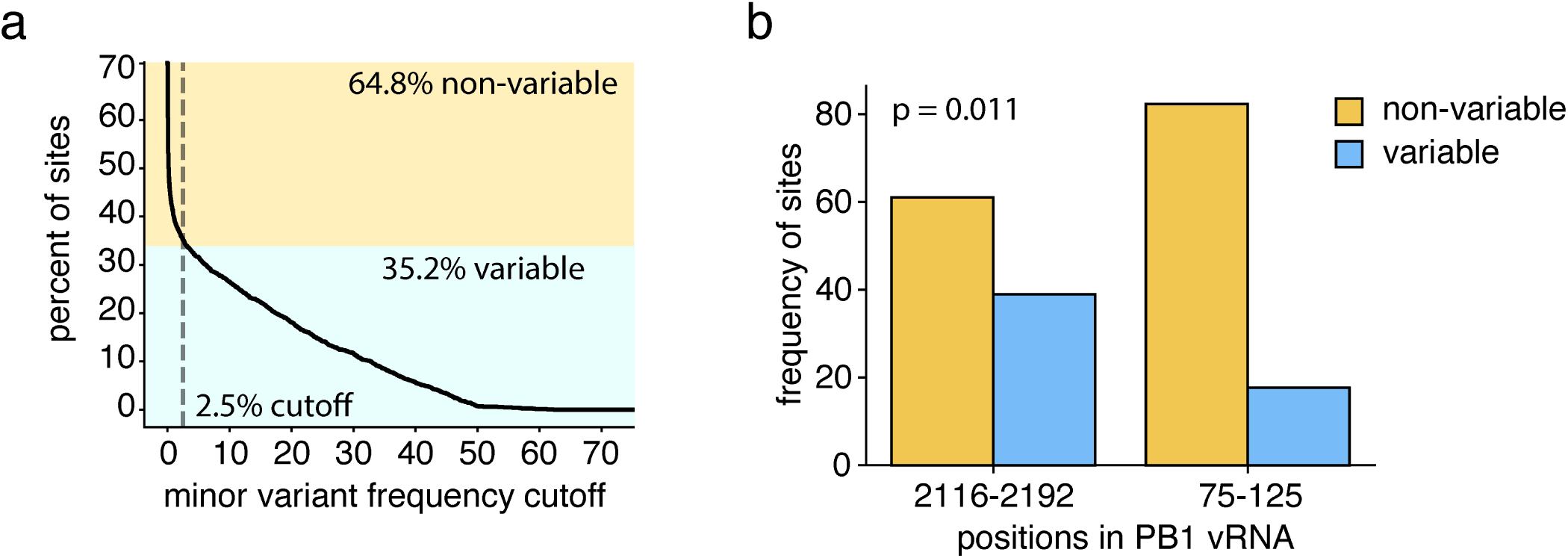
Conservation of regions from Fig. 4c across natural PB1 sequences. To compare natural diveristy, we procured PB1 sequences from the NCBI flu database on July 25th, 2024 that met the following prerequisites: full-length, 2341nt long, and exclude vaccine and laboratory strains. These sequences were then clustered by CD-HIT to an identity of 99%, and a single example of each cluster retained. Thereafter, sequences with ambiguities or indels relative to other PB1 sequences, were manually removed. This rsulted in 1939, high-quality, PB1 sequences that exhibit good alignment when simply lined-up with one-another, giving high confidence in the comparability of any given position, and, given the CD-HIT clustering, are unlikely to be aggressively weighted towards any single outbreak. **a** To define positions as variable or non-variable, we calculated the fraction of variable sites depending on the minimum minor variant frequency at which we consider a position variable. For instance, at the chosen cutoff, 2.5%, a position is considered variable if at least 2.5% of sequences do not match the major variant at that site. This cutoff was chosen to match the inflection point on our curve. **b** Frequency of variable sites within the given regions of PB1 as defined in 4c. Significant difference tested using Fisher’s exact test, p value shown.

**Supplementary Figure 4.**
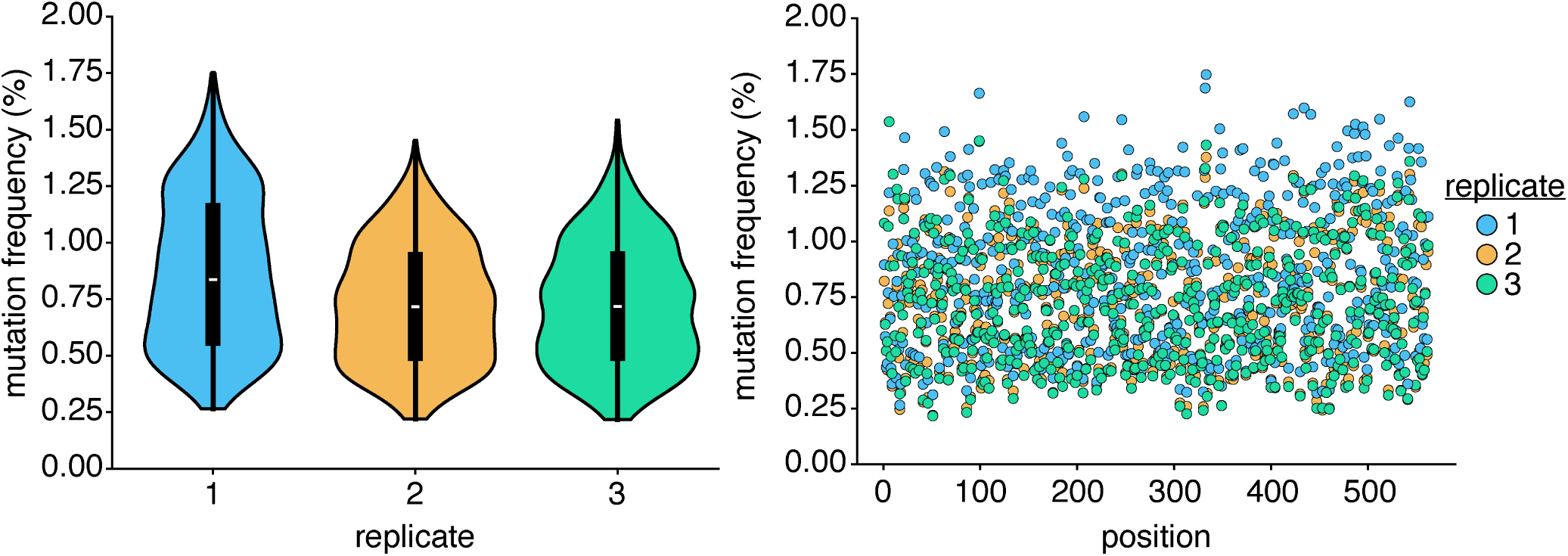
Generation of single-nucleotide variant libraries in PB1_177:385_. (left) Per site Mutation frequency as measured by Illumina sequencing of PB1_177:385_ libraries generated by mutagenic PCR across each of the three replicate libraries. Libraries were relatively uniform, with a median mutation frequency ranging from 0.716% to 0.836% (right) Data from (left) displayed per individual site in PB1_177:385_. Mutation rate was relatively uniform across this template, with no particular regions exhibiting particularly aberrant frequencies across three replicates.

**Supplementary Figure 5.**
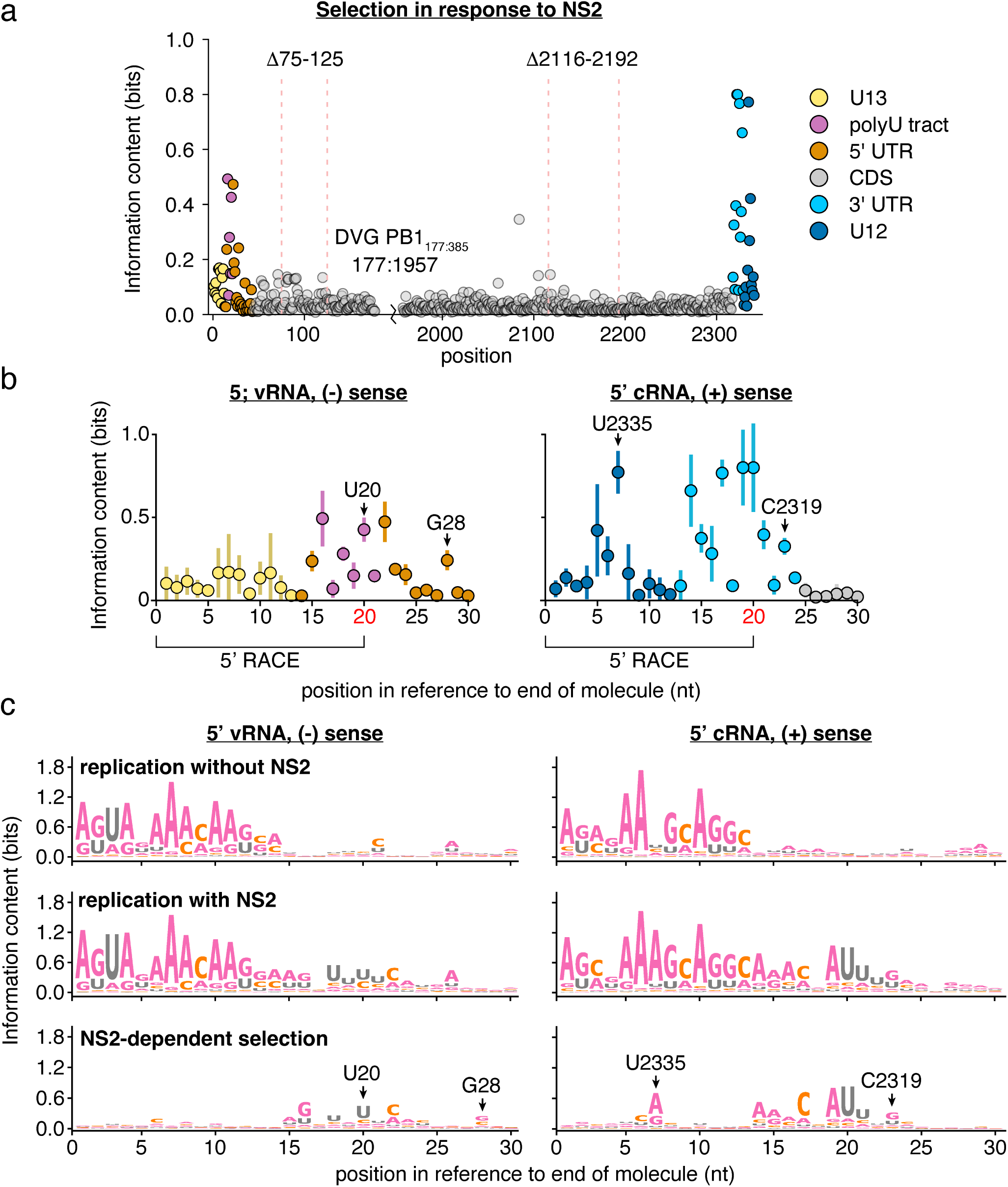
Information content of single-nucleotide variant libraries in PB1_177:385_. Information content calculated using Shannon entropy of the selection preferences at each position. Sequence logos generated by assigning this information content to each nucleotide in order of the strength of selection. **a** Information content analysis of data as presented in Fig. 5a. Each point represents the average of three replicates. **b** Information content analysis of data as presented in Fig. 5b, for NS2-dependent selection only. Mean and standard deviation displayed, n=3. Positions chosen for further analysis in Fig. 6 noted. **c** Sequence logo plots for positions displayed in Fig 5b. Data presented are the median values across three replicates. Positions chosen for further analysis in Fig. 6 noted. Inter-replicate correlation plots presented in (Supplementary Fig. 6)

**Supplementary Figure 6.**
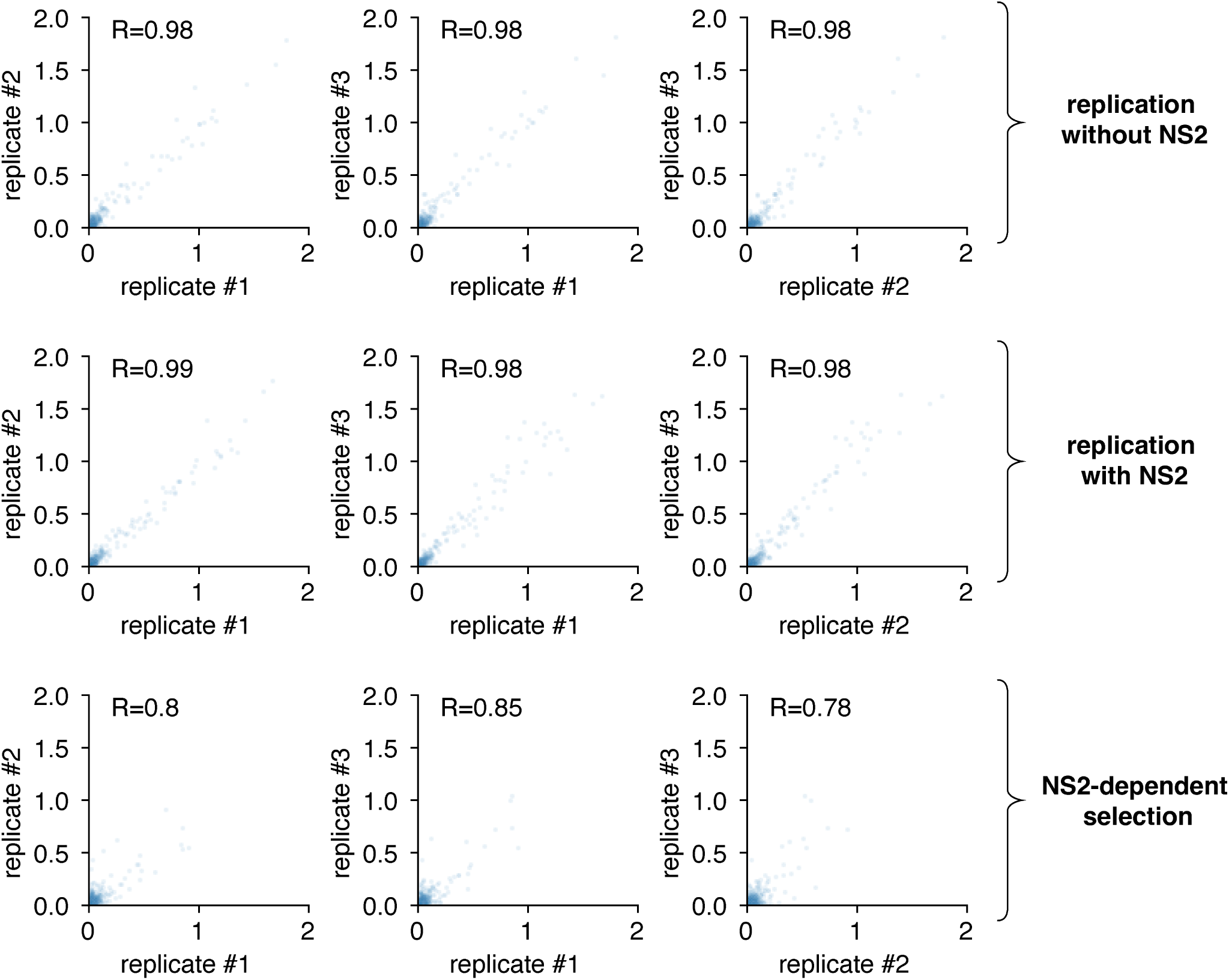
Inter-replicate correlation from Supplementary Fig. 5. Inter-replicate information content (total) values as calculated in Supplementary Fig 5. Values are in bits. R is the Pearson correlation coefficient.

**Supplementary Figure 7.**
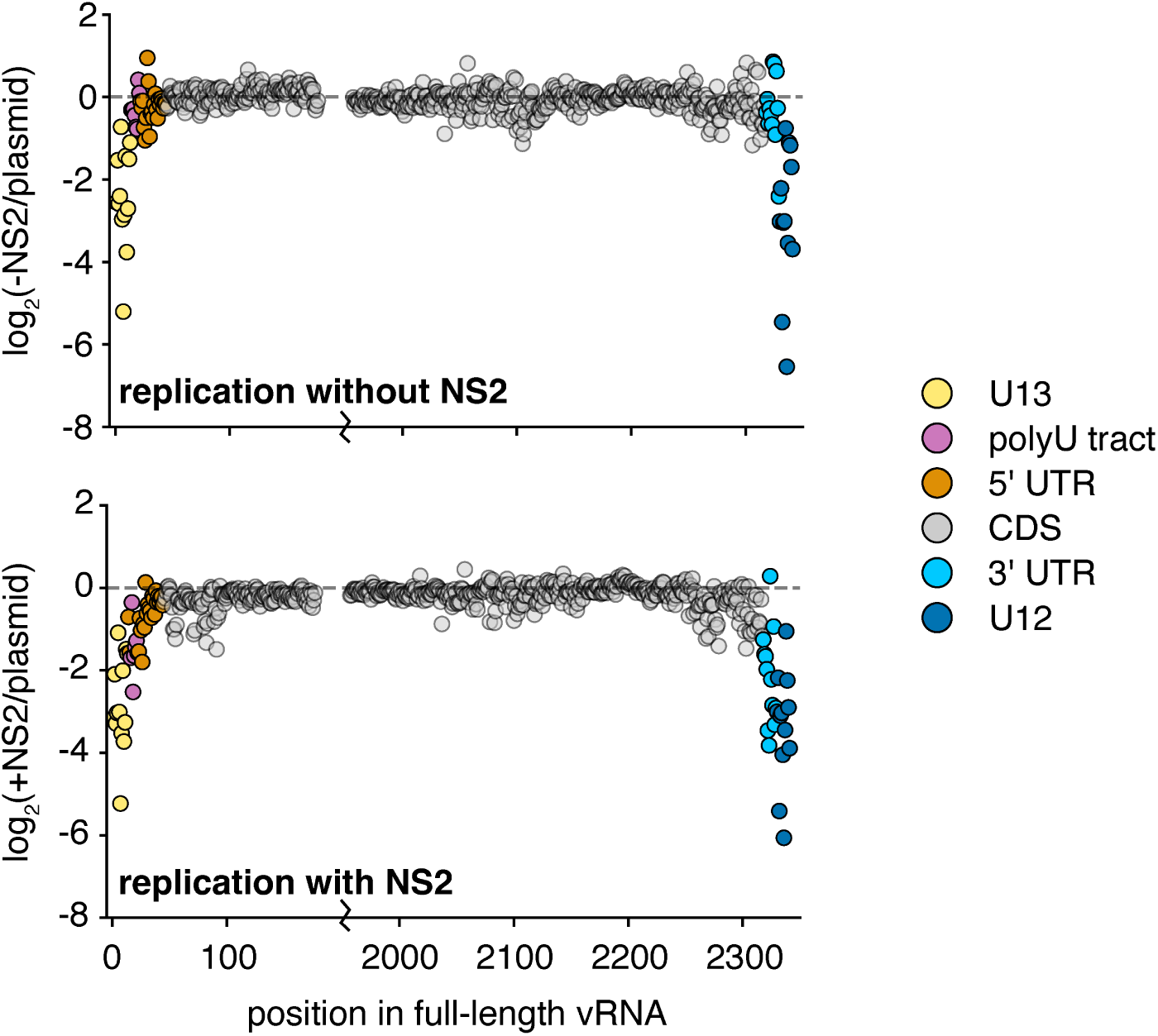
Full-length selection of subsets shown in Fig. 5b. Selection measured under each condition across all of PB1_177:385_ against non-wild-type nucleotides, average value across all three replicates provided.

**Supplementary Figure 8.**
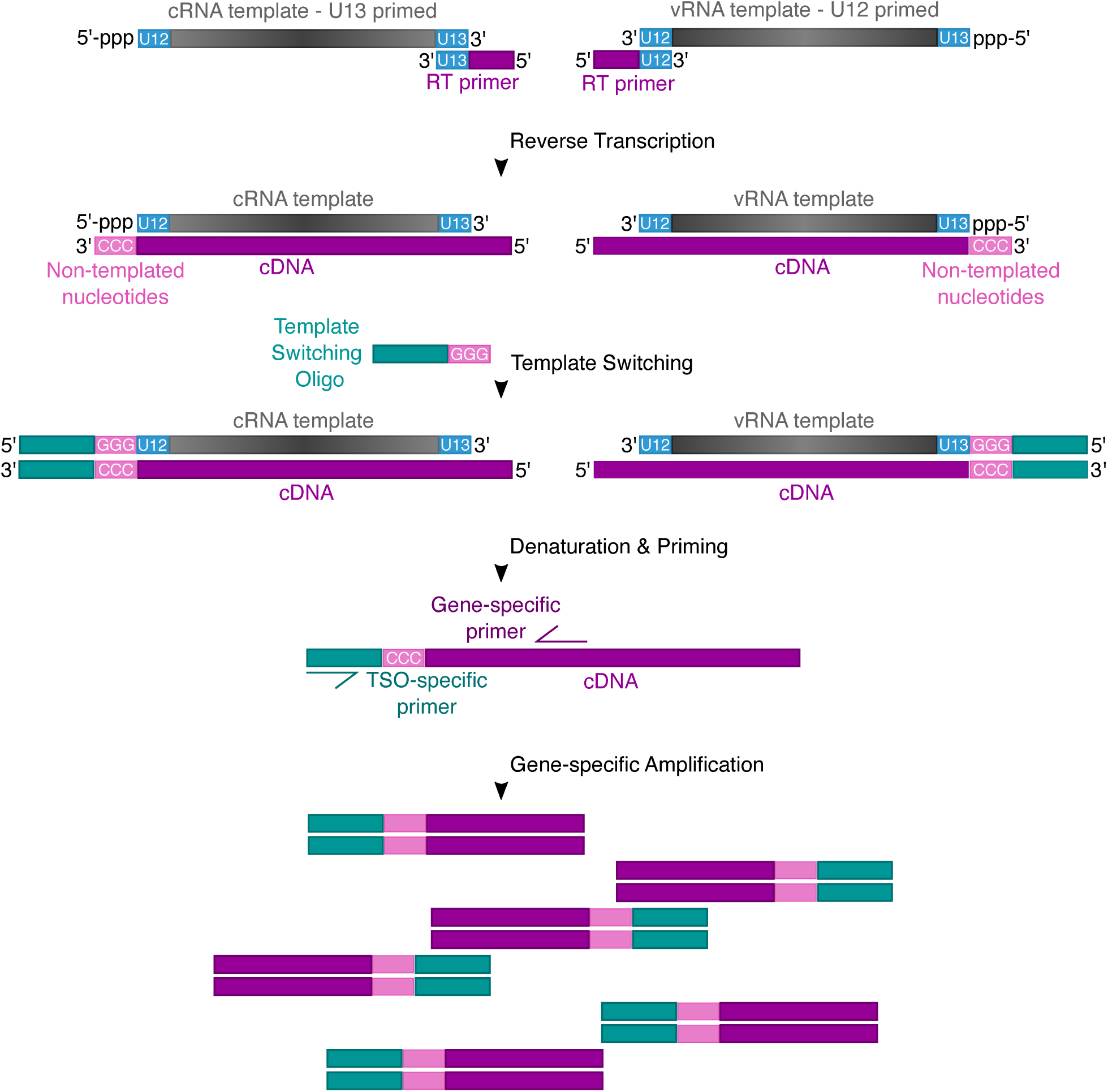
RACE inference of 5’ sequence. Schematic depicting the process by which 5’ sequence of cRNA and vRNA were measured.

**Supplementary Figure 9.**
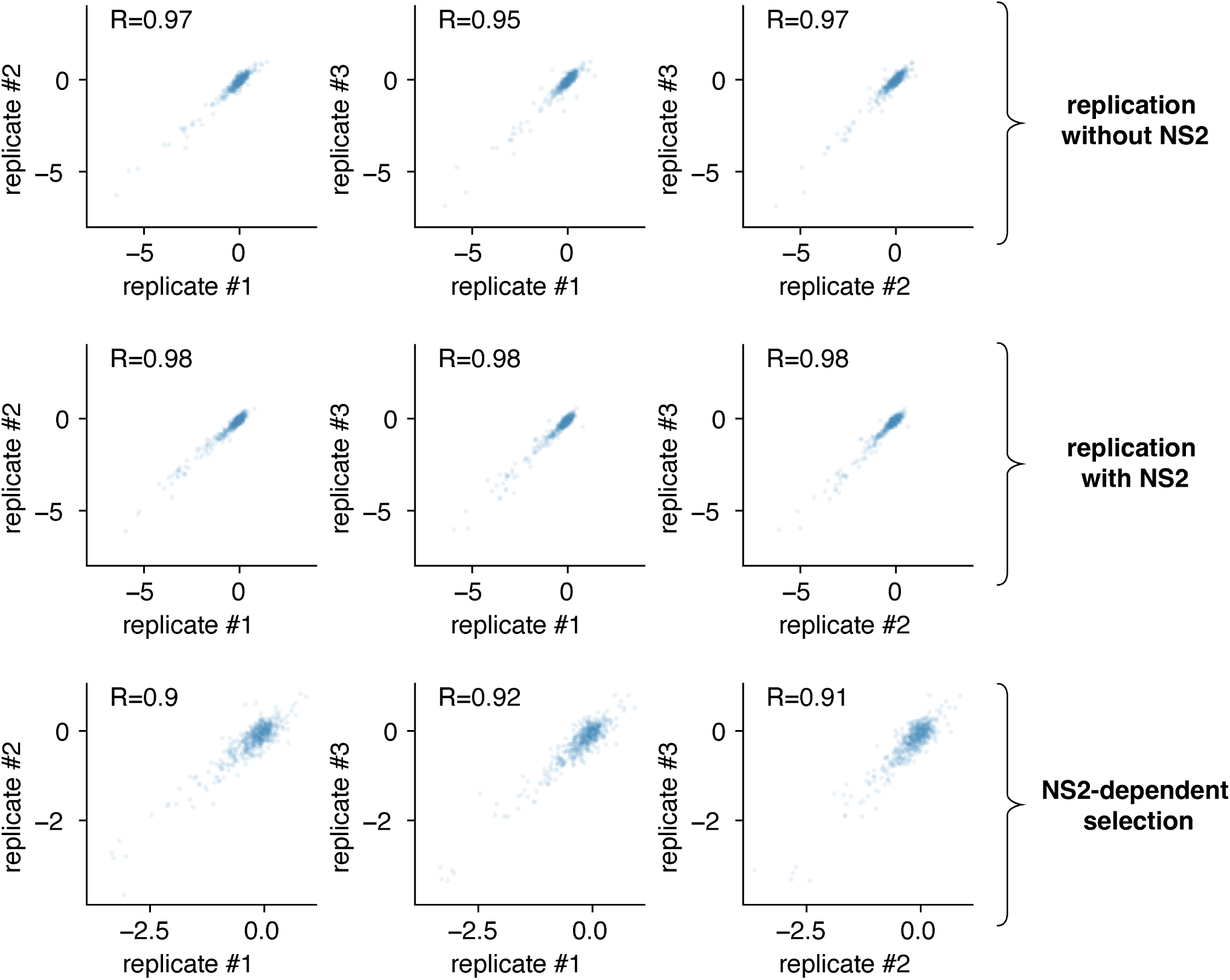
Inter-replicate correlation from Fig. 5. Inter-replicate selection on non-wild-type nucleotides as calculated in Fig 5. Values are in log_2_. R is the Pearson correlation coefficient.

**Supplementary Figure 10.**
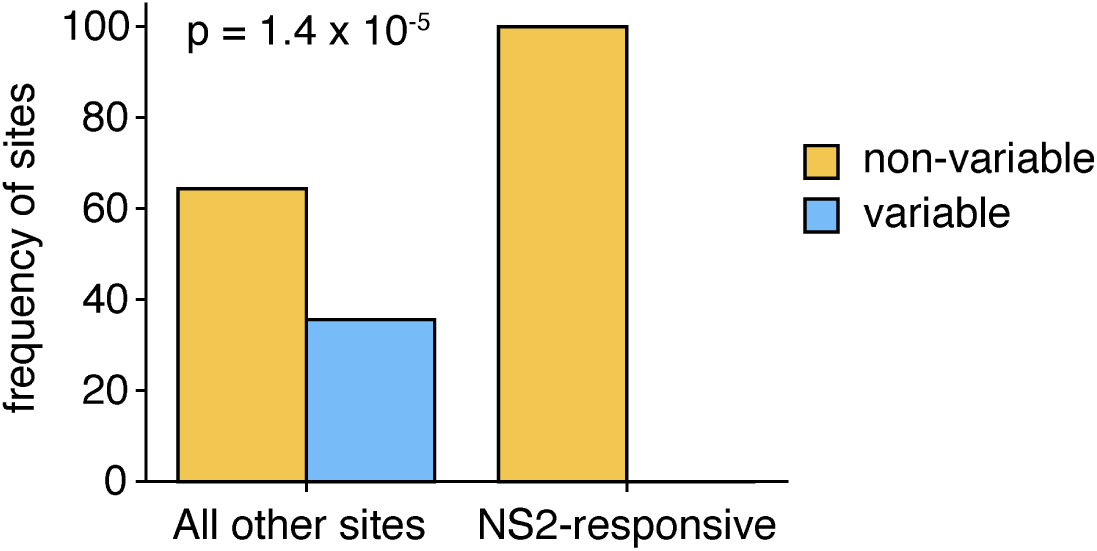
Conservation of sites identified in Fig. 5a as NS2-responsive across natural PB1 sequences. Analysis performed as in Supplementary Fig. 3 comparing the 27 sites we identify as critical to replication in the presence of NS2 as compared to all other sites in PB1. These sites are listed in Supplementary Table 1. Significant difference tested using Fisher’s exact test, p value shown.

**Supplementary Figure 11.**
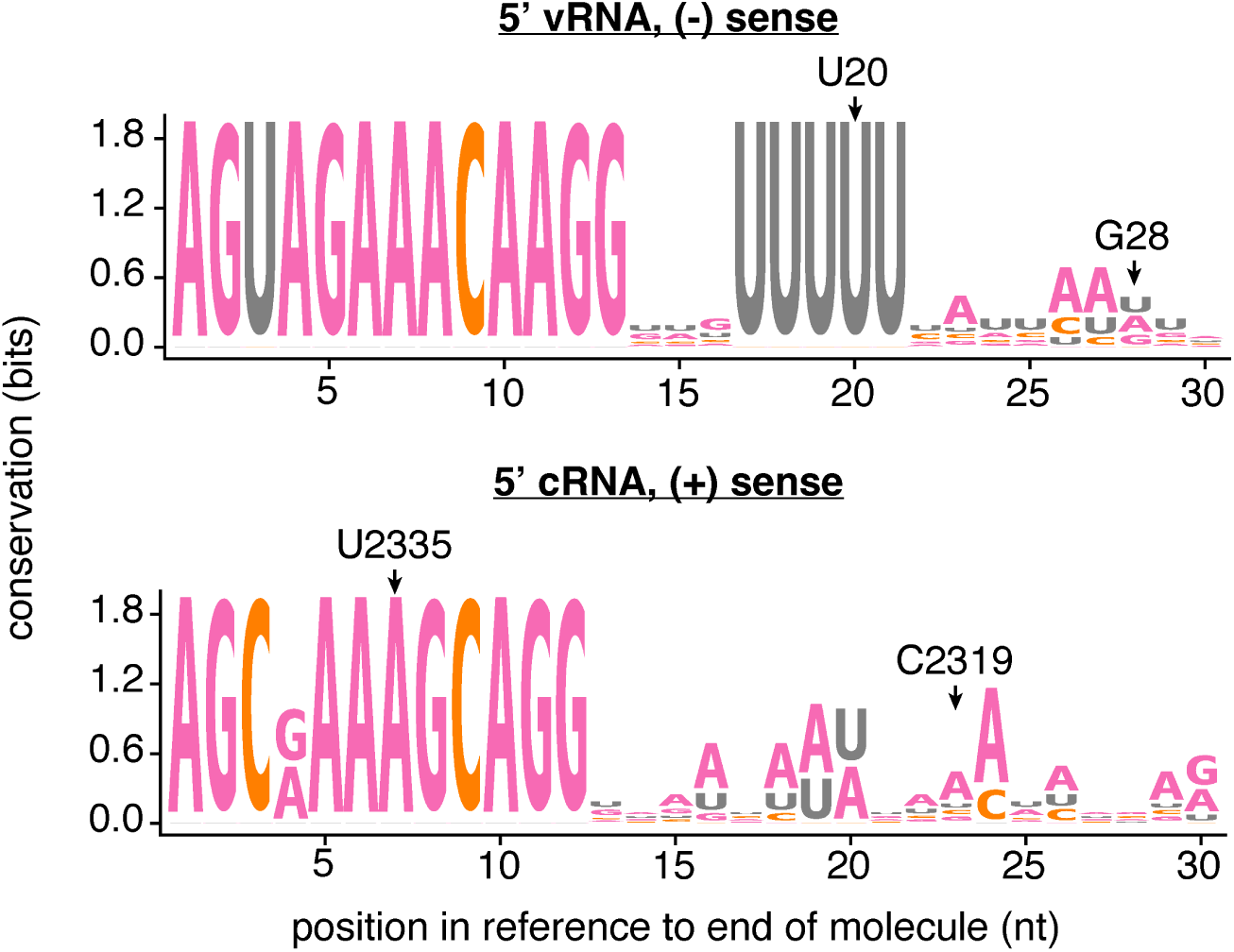
NS2 responsive sites are not identical between IAV segments. The first 30nt of the vRNA and cRNA were compared between the eight IAV segments, and sequence logo plots generated. The sites we tested in Fig. 6 are highlighted.

**Supplementary Figure 12.**
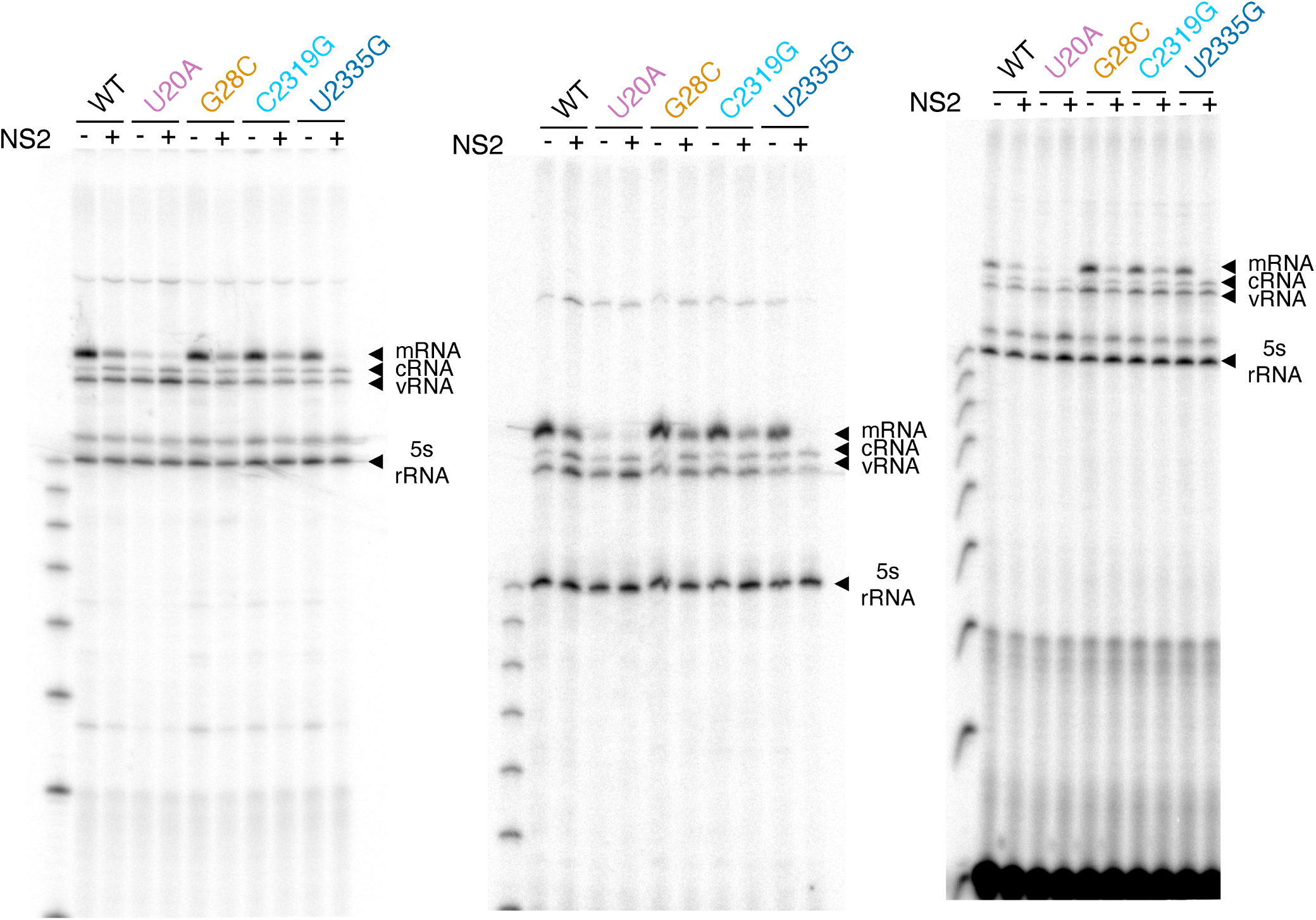
Primer extension analysis of various PB1_177:385_ variants in minimal replication assays with, and without, NS2. Full gels from primer extension analysis presented in Fig. 6b,c. First gel is the gel presented in Fig. 6b.

**Supplementary Figure 13.**
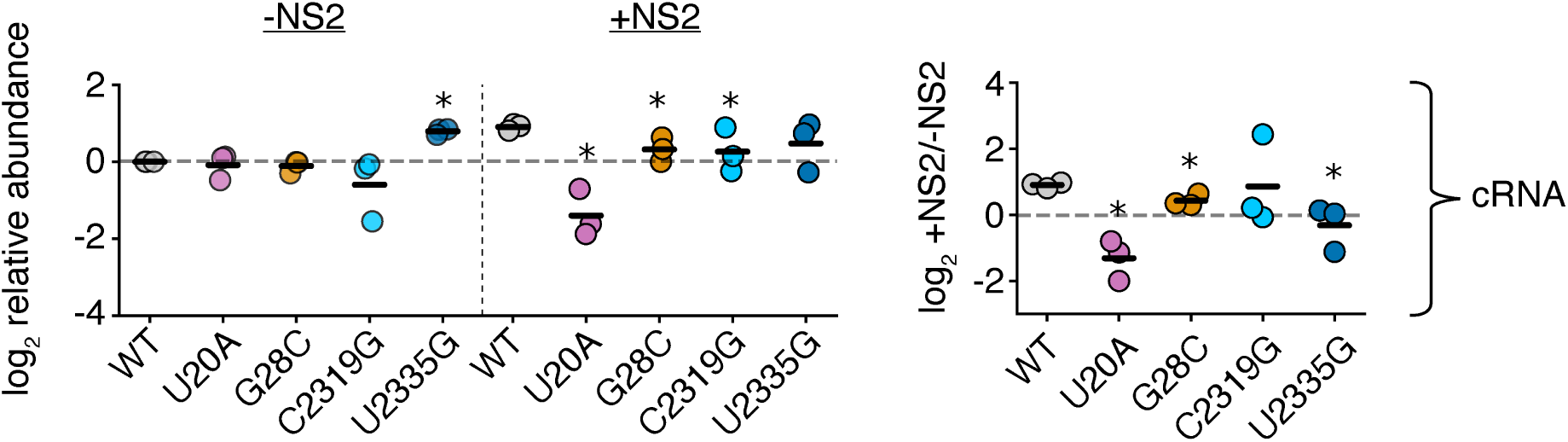
Mutations to NS2-dependent sites can impact replication of cRNA in full-length PB1. For all experiments, the canonical start codon was removed by mutagenesis to remove effects of additional expression of PB1. Quantitative analysis of primer extension as presented in Supplementary Fig. 14. All values in left two columns corrected against a parental template in the absence of NS2. Dotted line represents that value, points above indicate an increase in that molecular species, below, decrease. Values in the right column represent the ratio of points between the left two columns. Asterisks indicate values that are significantly decreased relative to the parental template, one-tailed t-test with Benjamini-Hochberg corrected FDR *<*0.1. Full gels presented in Supplementary Fig. 14. Individual replicates and mean presented, n=3.

**Supplementary Figure 14.**
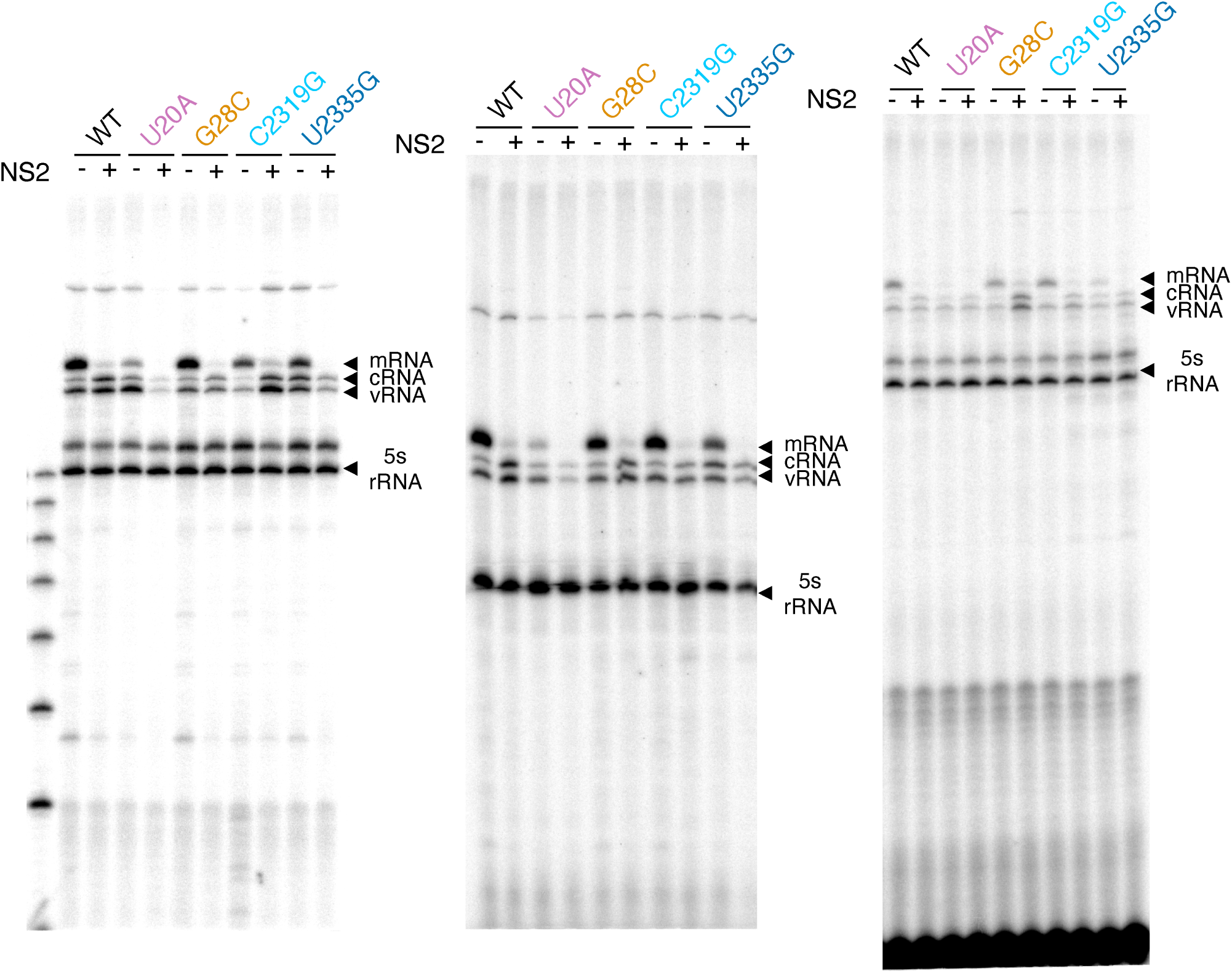
Primer extension analysis of various full-length PB1 variants in minimal replication assays with, and without, NS2. Full gels from primer extension analysis presented in Supplementary Fig. 13.

**Supplementary Table 1.** NS2-responsive sites. Table of all sites identified as meeting significance from Fig 5a.

| context | Nucleotide position (vRNA) | log2 NS2-dependent effect | pval | qval |
| --- | --- | --- | --- | --- |
| 5' UTR | 15 | -1.26642623 | 0.0053062 | 0.05516478 |
| polyU | 16 | -1.407887837 | 0.00580593 | 0.05530393 |
| polyU | 18 | -1.795893957 | 0.00826029 | 0.06727946 |
| polyU | 20 | -1.844711513 | 0.00254884 | 0.04325913 |
| polyU | 21 | -1.37795038 | 0.00446839 | 0.05022535 |
| 5' UTR | 22 | -1.462633616 | 0.02373645 | 0.09737143 |
| 5' UTR | 23 | -1.285950966 | 0.01276786 | 0.07827812 |
| 5' UTR | 28 | -1.90659152 | 0.0048643 | 0.05257184 |
| CDS | 55 | -1.101661611 | 0.00234029 | 0.04286222 |
| CDS | 56 | -1.100544239 | 0.00189633 | 0.04232093 |
| CDS | 71 | -1.326675414 | 0.01338237 | 0.07827812 |
| CDS | 72 | -1.03733307 | 0.0000459 | 0.01290355 |
| CDS | 82 | -1.178798373 | 0.0111679 | 0.07659062 |
| CDS | 83 | -1.067901354 | 0.00338681 | 0.04531879 |
| CDS | 91 | -1.372475945 | 0.00471253 | 0.05193021 |
| 3' UTR | 2319 | -1.555944694 | 0.00131708 | 0.03949276 |
| 3' UTR | 2320 | -1.015444989 | 0.02231737 | 0.09582604 |
| 3' UTR | 2321 | -1.696175204 | 0.0130107 | 0.07827812 |
| 3' UTR | 2322 | -3.011824531 | 0.0084403 | 0.06776352 |
| 3' UTR | 2323 | -3.149977188 | 0.00244055 | 0.04286222 |
| 3' UTR | 2325 | -3.02907342 | 0.00132183 | 0.03949276 |
| 3' UTR | 2326 | -1.928677286 | 0.02540826 | 0.09776512 |
| 3' UTR | 2327 | -1.56234586 | 0.00266523 | 0.04325913 |
| 3' UTR | 2328 | -3.048000255 | 0.00326677 | 0.04477866 |
| U12 | 2335 | -3.286629043 | 0.00325948 | 0.04477866 |
| U12 | 2339 | -1.072209116 | 0.0008218 | 0.03949276 |
| U12 | 2340 | -1.19881869 | 0.01855345 | 0.09035903 |

**Supplementary Data 1. Genome sequence of A/WSN/1933.**

Genome sequence of A/WSN/1933 used in this study.

**Supplementary Data 2. PB1_177:385_ sequence.**

Sequence of PB1_177:385_ from this study.

**Supplementary Data 3. NS2 sequences.**

NS2 sequences used in this study.

**Supplementary Data 4. Primer list.**

List of all primer sequences used in this study.

**Supplementary Data 5. PB1 sequences.**

PB1 sequences used to assess natural diversity.

